# Integrative analysis of transcription factor occupancy at enhancers and disease risk loci in noncoding genomic regions

**DOI:** 10.1101/262899

**Authors:** Shinya Oki, Tazro Ohta, Go Shioi, Hideki Hatanaka, Osamu Ogasawara, Yoshihiro Okuda, Hideya Kawaji, Ryo Nakaki, Jun Sese, Chikara Meno

## Abstract

Noncoding regions of the human genome possess enhancer activity and harbor risk loci for heritable diseases. Whereas the binding profiles of multiple transcription factors (TFs) have been investigated, integrative analysis with the large body of public data available so as to provide an overview of the function of such noncoding regions has remained a challenge. Here we have fully integrated public ChIP-seq and DNase-seq data (*n* ~ 70,000), including those for 743 human transcription factors (TFs) with 97 million binding sites, and have devised a data-mining platform —designated ChIP-Atlas—to identify significant TF-genome, TF-gene, and TF-TF interactions. Using this platform, we found that TFs enriched at macrophage or T-cell enhancers also accumulated around risk loci for autoimmune diseases, whereas those enriched at hepatocyte or macrophage enhancers were preferentially detected at loci associated with HDL-cholesterol levels. Of note, we identified “hotspots” around such risk loci that accumulated multiple TFs and are therefore candidates for causal variants. Integrative analysis of public chromatin-profiling data is thus able to identify TFs and tissues associated with heritable disorders.

## INTRODUCTION

The vast majority of noncoding regions in the human genome is thought to possess biological activity (ENCODE Project Consortium 2012). These regions include enhancers, the activity of which is regulated by the binding of transcription factors (TFs). Mutations in TF genes or TF binding sites thus give rise to disorders such as cancer and autoimmune diseases, whereas forced expression of TFs is able to switch or reprogram cell fates (Chambers and Studer 2011; Lin et al. 2013; Ward and Kellis 2012b; Gutierrez-Arcelus et al. 2016; Matharu et al. 2015). Genome-wide association studies (GWASs) have also underscored the importance of noncoding regions with the finding that most single nucleotide polymorphisms (SNPs) associated with heritable disorders are distributed within such genomic regions—and, in particular, tend to be located at DNase I–hypersensitive sites (DHSs) or TF binding sites (ENCODE Project Consortium 2012; Maurano et al. 2012; Schaub et al. 2012). Characterization of the pattern of TF binding to enhancer regions is therefore fundamental to an understanding of the physiological relevance of noncoding regions to cell type specification and human health.

Genome-wide chromatin-profiling approaches such as chromatin immunoprecipitation–sequencing (ChIP-seq) and DNase-seq have been applied to annotate TF binding sites and DHSs, respectively. Integrated chromatin-profiling data have been compiled by the ENCODE consortium (ENCODE Project Consortium 2012) and have served as a global resource for exploration of TFs enriched at tissue-specific enhancers and at SNPs identified by GWASs (Thurman et al. 2012; Xie et al. 2013; Griffon et al. 2015; Schaub et al. 2012; Ng et al. 2014; Lindström et al. 2014; Farh et al. 2014). In addition, several tools have been developed to perform such enrichment analysis and functional annotation in a manner largely dependent on the ENCODE project data (Guo et al. 2015; Coetzee et al. 2012; Guo et al. 2014; Boyle et al. 2012; Ward and Kellis 2012a). On the other hand, a substantial amount of chromatin-profiling data has been presented by various smaller projects (**Fig. 1A**). Although these latter data are publicly available from Sequence Read Archive (SRA) of NCBI (https://www.ncbi.nlm.nih.gov/sra), they have been made use of to a lesser extent by the research community than have the ENCODE data for several reasons: (1) unlike the ENCODE data, only the raw sequence data are archived in most cases, necessitating extensive bioinformatics analysis; (2) metadata such as antigen and cell type names are often ambiguous as a result of the use of orthographic variants such as abbreviations and synonyms; and (3) integrative analysis of such data requires sophisticated skills for data-mining and abundant computational resources.

**Figure 1.**
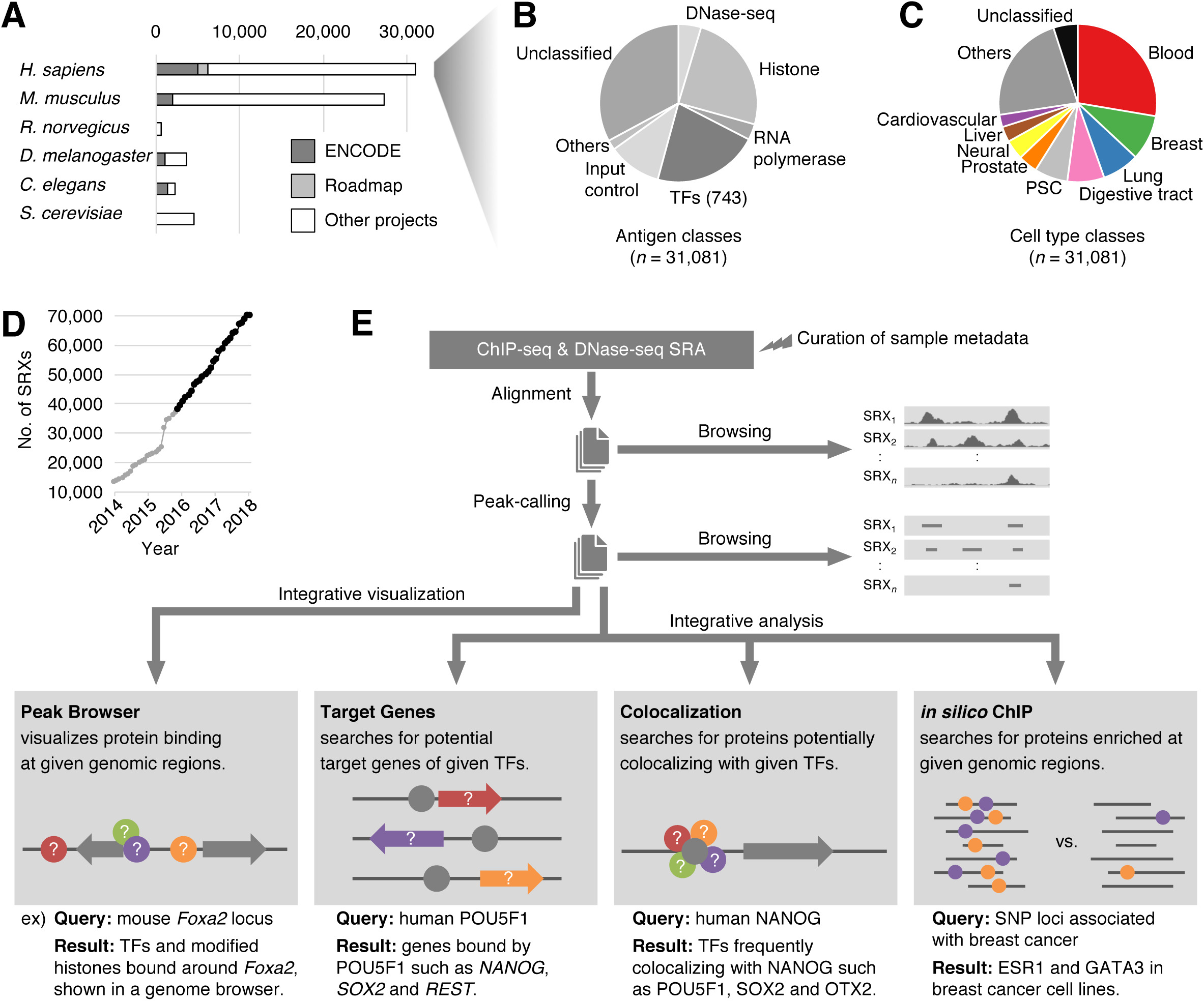
Overview of the ChIP-Atlas data set and computational processing. (*A*) Numbers of ChIP-seq and DNase-seq experiments recorded in ChIP-Atlas (as of February 2018), indicating the proportion of the data for each species derived from ENCODE, Roadmap Epigenomics, and other projects. (*B, C*) Numbers of experiments according to antigen (*B*) or cell type (*C*) classes for human data. PSC, pluripotent stem cell. (*D*) Cumulative number of SRX-based experiments recorded in ChIP-Atlas. Data published before and after the launch of ChIP-Atlas in December 2015 are shown in gray and black, respectively. (*E*) Overview of data processing. Raw sequence data are downloaded from NCBI SRA, aligned to a reference genome, and subjected to peak-calling, all of which can be monitored with the genome browser IGV. All peak-call data are then integrated for browsing via the “Peak Browser” function, and they can be analyzed for TF-gene (“Target Genes”) or TF-TF (“Colocalization”) interactions as well as subjected to enrichment analysis (“*in silico* ChIP”). All of the results are tagged with curated sample metadata such as antigen and cell type names. In the diagrams, gray components (circles, TFs; arrows, genes) indicate queries by the user, with colored components representing the returned results.

In an attempt to provide a global understanding of the TF landscape in noncoding regions of the human genome based on all chromatin-profiling data in the public domain, we have now developed a database and associated data-mining tool that we have designated ChIP-Atlas (http://chip-atlas.org). Taking advantage of the full data set, we explored TFs enriched at tissue-specific enhancers and around disease-associated noncoding SNPs. The unprecedented scale of the assembled chromatin-profiling data and supported integrative analyses has provided insight into the TF landscape at noncoding regions and thereby shed light on mechanisms of cell type– specific enhancer activation and pathological processes underlying heritable disorders.

## RESULTS

### Overview of the data set and design of analyses

ChIP-Atlas collects and integrates almost all of the public ChIP-seq and DNase-seq data for six organisms (90% of experiments for all organisms) (**Fig. 1A**–**C**). The source data are derived from NCBI SRA, in which each experiment is assigned an ID with a prefix of SRX, DRX, or ERX (hereafter collectively referred to as SRXs), and they have been updated monthly since the launch of the project in December 2015 (**Fig. 1D**). We manually curate the names of antigens and cell types according to commonly or officially adopted nomenclature. The antigens and cell types are further sorted into “antigen classes” and “cell type classes,” allowing categorization and extraction of data for given classes (**Fig. 1B,C**; **Supplemental Fig. S1A)**. The sequence data are aligned to a reference genome and subjected to peak-calling, and the results are readily downloaded and browsed in the genome browser IGV (Robinson et al. 2011) (**Fig. 1E**). ChIP-Atlas is superior to other similar services (Sun et al. 2013; Yevshin et al. 2017; Yang et al. 2013) in terms of both the amount of data (69,392 SRXs as of February 2018) and the number of covered organisms, as represented in a recent report comparing such services (Yevshin et al. 2017). Furthermore, notable features unique to ChIP-Atlas are that we monthly integrate whole peak-call data for browsing with IGV and analyze them integratively not only to reveal TF-gene and TF-TF interactions but also to allow enrichment analysis for given genomic intervals based on global protein-genome binding data—as reflected in the four functions shown in **Fig. 1E**.

All peak-call data recorded in ChIP-Atlas can be graphically displayed with the “Peak Browser” function at any genomic regions of interest (ROIs). To implement this function, we integrated a large amount of peak-call data (for example, 478 million and 298 million peaks for the human and mouse genomes, respectively), indexed them for IGV, and constructed a Web interface that externally controls IGV preinstalled on the user’s machine (Mac, Windows, or Linux platforms). For instance, on specification of ChIP-seq data for mouse TFs on the Web page (**Supplemental Fig. S1B**), the corresponding results are streamed into IGV as shown in **Fig. 2**, suggesting that the mouse *Foxa2* gene promoter is bound by multiple TFs in the liver (**Fig. 2**, center), that expression of the gene is suppressed by Polycomb group 2 proteins such as Suz12 and Ezh2 in embryonic stem cells (ESCs) (**Fig. 2**, left), and that the upstream region of *Foxa2* may possess insulator activity due to Ctcf binding in multiple cell types (**Fig. 2**, right). The colors of peaks indicate the statistical significance values calculated by the peak-caller MACS2 (MACS2 scores). Clicking on a peak opens a Web page containing detailed information including sample metadata, library description, and read quality (**Supplemental Fig. S1C**) as well as controllers to display the alignment data in IGV (**Fig. 2**, top). ChIP-Atlas thus allows not only visualization of the data for each experiment but also browsing of an integrative landscape of multiple chromatin-profiling results, potentially providing insight into the location of functional regions (enhancers, promoters, and insulators) and the corresponding regulatory factors (TFs and histone modifications). Since the public release, ChIP-Atlas has been used all over the world (> 100,000 page views by ~ 3,000 unique users per month at the submission of this manuscript) and has already been cited by 25 papers to make full use of public ChIP-seq data (http://chip-atlas.org/publications). Taking advantage of the large amount of TF binding data, we explored TFs enriched at given genomic features such as enhancers and GWASs-identified SNP loci.

**Figure 2.**
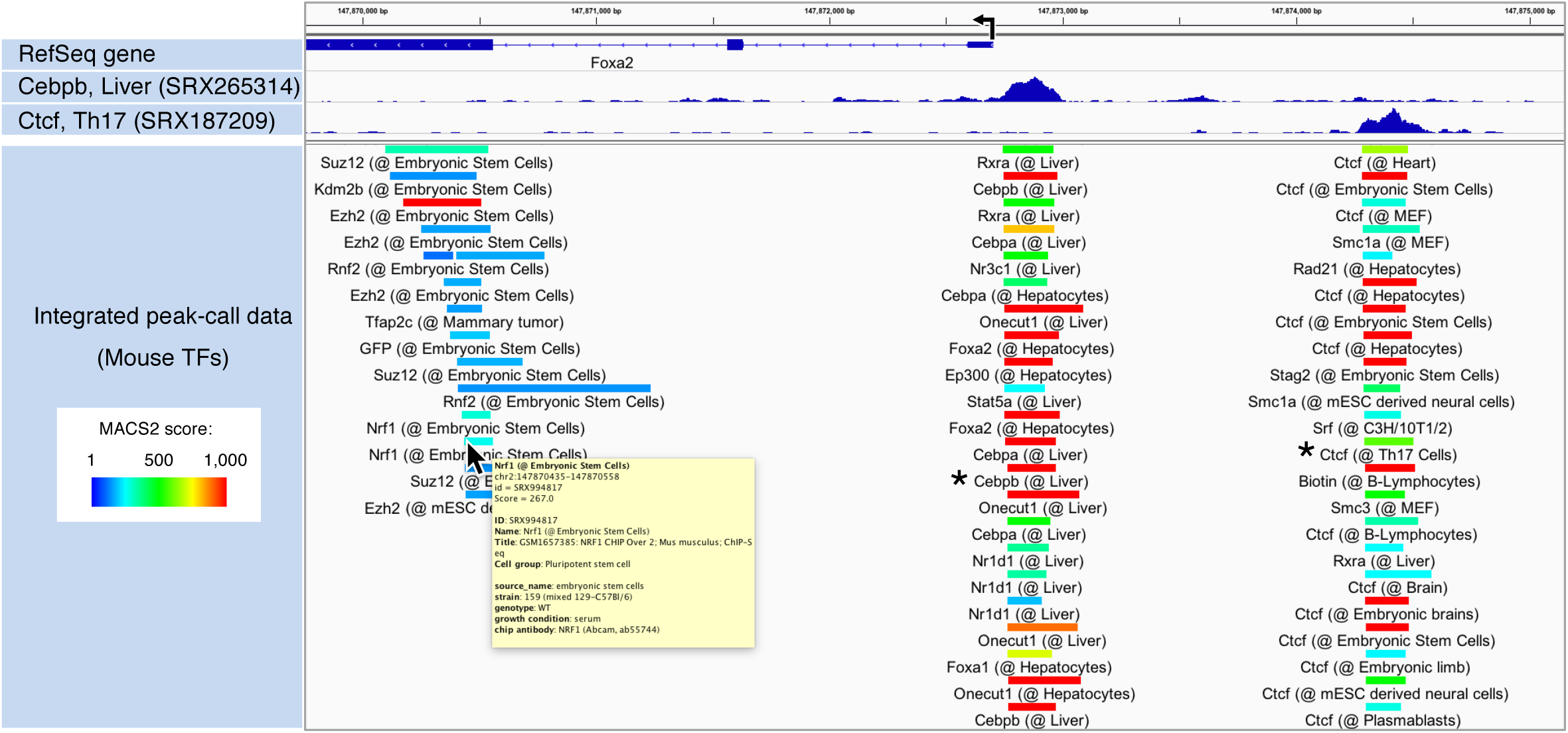
Example of processed data visualized with “Peak Browser” of ChIP-Atlas. ChIP-Atlas peak-call data for TFs around the mouse *Foxa2* locus are shown in the IGV genome browser for settings of the “Peak Browser” Web page shown in **Supplementary Figure S1B**. Bars represent the peak regions, with the curated names of the antigens and cell types being shown below the bars and their color indicating the ^score calculated with the peak-caller MACS2 (–log10[*Q* value]). Detailed sample^ information (yellow window) appears on placing the cursor over each bar. Clicking on the bars (asterisks) enables display of the alignment data (top) and detailed information about the experiments (**Supplementary Fig. S1C**).

### Dissection of TF-Dependent Regulation of Tissue-Specific Enhancer Activity

Although substantial progress has been made in the identification of enhancers activated in specific cell types (Andersson et al. 2014; ENCODE Project Consortium 2012), the TFs that coordinate regulation at these loci are less well characterized. To address this deficiency, we searched for TFs that are enriched at enhancer regions activated in a tissue-specific manner with the use of “*in silico* ChIP” of ChIP-Atlas, a unique tool to search for TFs enriched at a batch of genomic ROIs. On submission of two sets of genomic regions (ROIs and background regions), this service evaluates all SRXs to count the overlaps between the peaks and submitted regions and then returns enrichment analysis data including SRX IDs, TFs, cell types, and *P* values (**Fig. 3A**; **Supplemental Fig. S2**). FANTOM5 “predefined enhancer data” served as the list of tissue-specific enhancers, which include 20,795 enhancers significantly activated in at least one cell type (*n* = 69) or anatomic facet (*n* = 41) (Andersson et al. 2014). As a proof of principle, we selected hepatocyte-specific enhancers as the ROIs (*n* = 286) and applied “*in silico* ChIP” to search for TFs enriched at these enhancers relative to all other enhancers (*n* = 20,509). Significantly enriched TFs included HNF4A/G and FOXA1/2 (*P* < 1 × 10^−21^) (**Fig. 3B,C**), which are required for liver development and are able to directly reprogram skin fibroblasts into hepatocyte-like cells (Du et al. 2014; Sekiya and Suzuki 2011; Smith et al. 2013). Furthermore, TFs for the top 15 ranked SRXs included SP1, RXRA, CEBPB, and JUND, all of which function in the liver-biliary system according to Mouse Genome Informatics (MGI) phenotype collections (MP:0005370). In addition, the predominant cell type from which enriched SRXs were derived was Hep G2 in the Liver class, even though the number of experiments in this class is relatively small for human (**Fig. 1C**), thus demonstrating the high detection power of our analysis.

**Figure 3.**
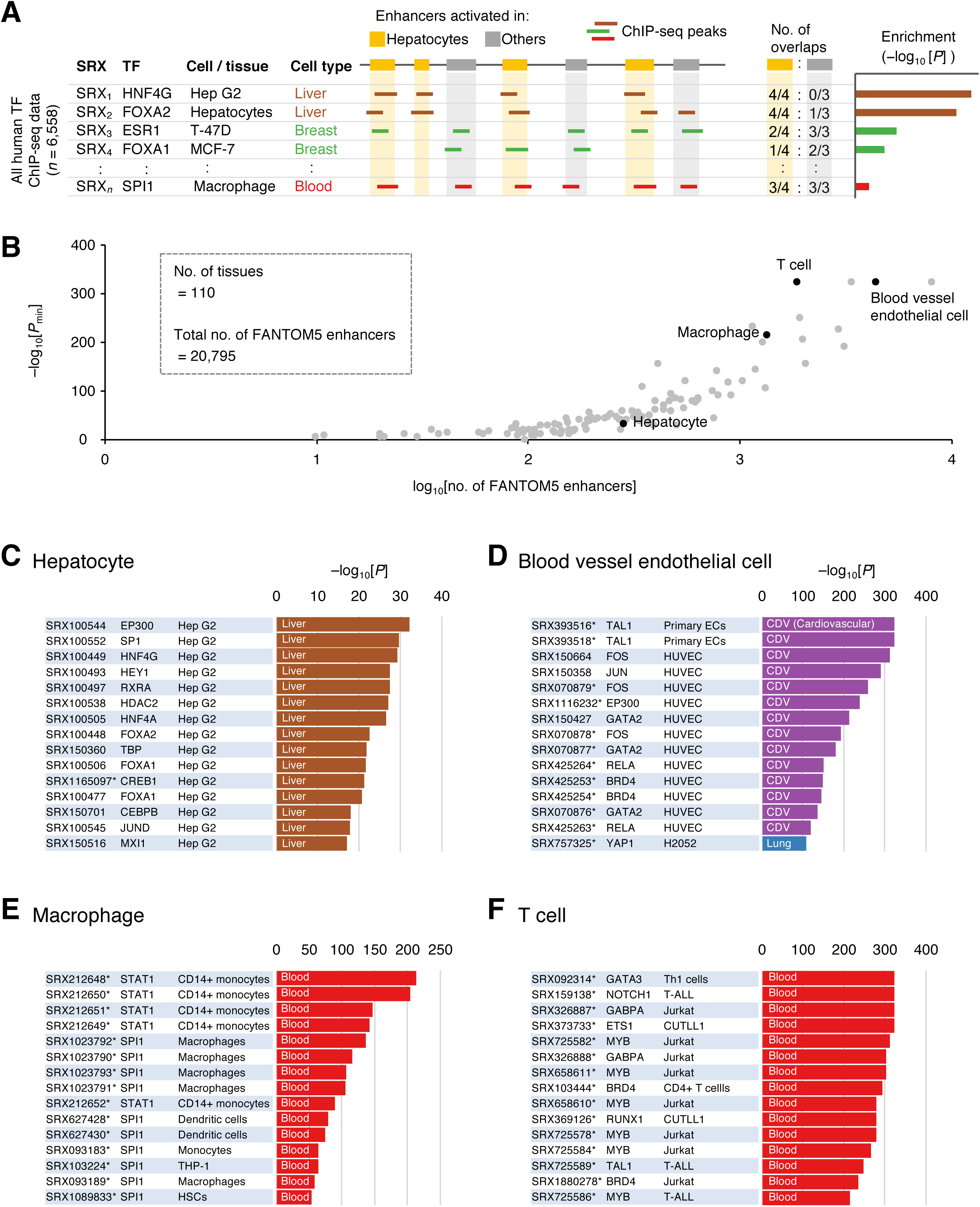
Analysis of TF enrichment at FANTOM5 tissue-specific enhancers with “*in silico* ChIP” of ChIP-Atlas. (*A*) Overview of enrichment analysis. On submission of BED-formatted genomic regions for hepatocyte enhancers (orange) or enhancers activated in other tissues (gray), a computational server counts the overlaps with the peaks of all SRXs (left). After evaluation of the significance of enrichment with Fisher’s exact test (right), the analyzed data are returned to the machine of the user as shown in **Supplementary Figure S2**. (*B*) Summarized results for TF enrichment analysis with “*in silico* ChIP” showing *P*_min_ and numbers of enhancers specifically activated in 110 tissues identified by the FANTOM5 project. (*C*–*F*) ChIP-seq data enriched for enhancers of hepatocytes (*C*), blood vessel endothelial cells (*D*), macrophages (*E*), or T cells (*F*) relative to all other FANTOM5 enhancers. The bar charts indicate *P* values for enrichment, with the colors indicating the cell types examined in the experiments according to the palette shown in **Fig. 1C**. Asterisks next to SRX IDs indicate that the ChIP-seq data originated from projects other than ENCODE or Roadmap Epigenomics.

We next examined all facets of FANTOM5 predefined tissue-specific enhancers (**Fig. 3B**, and see URL in **Supplemental Table S1** for the latest complete results). One of our most significant findings was obtained for enhancers specifically activated in blood vessel endothelial cells (**Fig. 3D**), with a minimum *P* value (*P*_min_) of <1 × 10^−324^. All cell types of the top 14 ranked SRXs were endothelial cells in the Cardiovascular class, and the enriched TFs included TAL1, JUN, EP300, GATA2, and YAP1, all of which contribute to blood vessel morphology phenotypes (MP:0001614). The number of ChIP-seq experiments in the Cardiovascular class is also small, demonstrating that “*in silico* ChIP” analyses have a low susceptibility to bias with regard to the number of data sets recorded in ChIP-Atlas.

Another remarkable finding was the pronounced enrichment of SRXs of the Blood class at enhancers of blood-related cells. In particular, SPI1 and STAT1 were enriched at macrophage enhancers (*P*_min_= 1 × 10^™214^), with the cell types being cells of the myeloid lineage including macrophages and their progenitors (monocytes) as well as dendritic cells and monocytic cell lines (THP-1) (**Fig. 3E**). Of note, SPI1 (also known as PU.1) is able to reprogram fibroblasts into macrophages (Feng et al. 2008), suggesting that pioneer activity of SPI1 serves to collectively activate macrophage-specific enhancers prior to macrophage development.

T cell enhancers also yielded a highly significant result (*P*_min_ < 1 × 10^−324^). Although the enriched cell type class was also Blood, other features were distinct from those of macrophage enhancers. The enriched cell types were thus of lymphoid origin, including cells of the T cell lineage, leukemia (T–acute lymphoblastic leukemia, or T-ALL), and lymphoid cell lines (Jurkat and CUTLL1). Furthermore, enriched TFs included GATA3, NOTCH1, GABPA, ETS1, MYB, and RUNX1, all of which are associated with T cell phenotypes (MP:0008037) (**Fig. 3F**), suggesting that these TFs might be as-yet-unidentified factors for direct reprograming to the T cell lineage.

Together, our enrichment analyses thus revealed TFs that show significant binding to tissue-specific enhancers, suggesting that these proteins may function as master regulators of tissue specificity and commitment to corresponding cell types. Furthermore, it should be noted that most of the enriched SRXs in these analyses originated from projects other than ENCODE or Roadmap Epigenomics (asterisks in **Fig. 3C**–**F**), highlighting the detection power conferred by the large amount of real binding data archived in ChIP-Atlas.

### Isolation of TFs enriched around noncoding SNPs identified by GWASs

GWASs have identified thousands of SNPs associated with human phenotypes and diseases. However, most SNPs associated with genetic diseases lie within noncoding regions (Maurano et al. 2012), with the result that the mechanisms by which they contribute to pathological processes remain unclear. Furthermore, causal variants do not necessarily coincide with marker SNPs identified by GWASs, but may instead reside within linkage disequilibrium (LD) blocks of such SNPs. Given that the loci of causal SNPs tend to have functional properties such as cis-regulatory activity characterized by DNase-hypersensitivity (Brown et al. 2017), we assumed that causal SNPs, rather than marker SNPs, would be coordinated by multiple TFs. To test this notion, we examined all cell types for DHSs located within LD blocks (*r*^2^ > 0.9) of noncoding marker SNPs (referred to as LD-DHSs) (**Supplemental Fig. S3**). We first tested the LD-DHSs associated with breast cancer (BRC) as ROIs, with those of other traits being selected as background regions, for “*in silico* ChIP” analysis. Of note, enrichment of ChIP-seq data was most significant (*P* = 2 × 10^−14^) for ESR1 (estrogen receptor 1) in the BT-474 BRC cell line (**Fig. 4A**,**B**), likely reflecting the fact that estrogen signaling contributes to the proliferation of BRC cells (Carroll et al. 2006, 2005). In addition, the top 15 SRXs included other BRC cell lines (T-47D, MCF-7, and ZR-75-1) and the TFs ARNTL, ESRRA, TFAP2C, GATA3, HIF1A, and NR3C1. Significant enrichment of these TFs was not detected by enrichment analysis based on ENCODE ChIP-seq data (Schaub et al. 2012) or by a motif survey (Maurano et al. 2012), suggesting that the large amount of real binding data probed by ChIP-Atlas contributed to the detection of these novel TFs. Importantly, we noticed that there were some “hotspots” among LD-DHSs where multiple TFs accumulated (**Supplemental Fig. S4**). As an example, the upstream region of *KLF4* contains two marker SNPs for BRC susceptibility (rs10759243 and rs865686), within the LD blocks of which we identified two hotspots occupied by TFs such as ESR1, TFAP2C, GATA3, and NR3C1 in BRC cell lines (**Fig. 4B**, bottom right). We surveyed the functional annotations for common SNPs at these hotspots and found that one such SNP (rs5899787) in the LD block of rs865686 has been shown to be causally associated with BRC by large-scale fine-mapping and reporter assays (Orr et al. 2015), suggesting that the hotspots we identified might be indicative of causal variants. On the other hand, we did not find any annotations for common SNPs (rs12350528 and rs1832880) within the rs10759243-asscociated hotspot (**Fig. 4B**, bottom left), which may thus be a previously unidentified causal variant for BRC.

**Figure 4.**
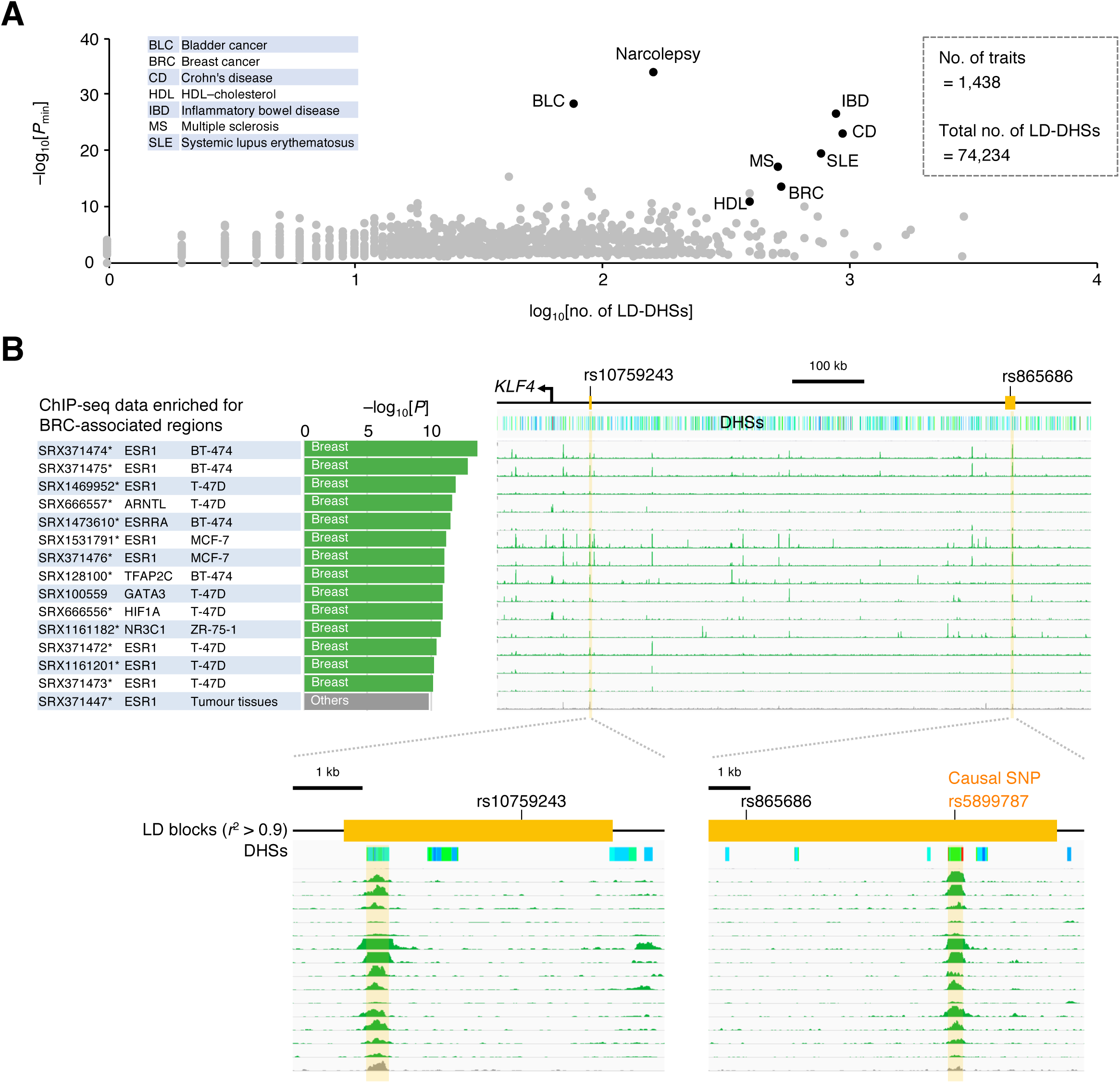
Analysis of TF enrichment around noncoding GWAS-identified SNPs with “*in silico* ChIP” of ChIP-Atlas. (*A*) Summarized results of TF enrichment analysis with ^“*in silico* ChIP” showing *P*^min ^and numbers of LD-DHSs for 1438 traits archived in the^ NHGRI-EBI GWAS catalog. (*B*) ChIP-seq data enriched for the regions associated with BRC. Alignment data are also shown around two marker SNPs associated with BRC, with the *y*-axes ranging from 0 to 1 RPM units.

We next performed enrichment analyses for all the traits archived in the NHGRI-EBI GWAS catalog (Welter et al. 2014) as we did for BRC (total of 74,234 LD-DHSs for 1438 traits) (**Fig. 4A**, and see URL in **Supplemental Table S1** for the latest complete results). One significant result was obtained for inflammatory bowel disease (IBD), for which the associated regions showed enrichment for ChIP-seq data derived from immune-related cell types and TFs (*P*_min_ = 3 × 10^−27^) (**Fig. 5A**). Although the ChIP-Atlas data set contains a substantial amount of ChIP-seq data derived from the digestive tract (**Fig. 1C**), none of the corresponding SRXs showed enrichment in this analysis, consistent with evidence that an autoimmune response is the primary cause of IBD. Such enrichment for blood-related cell types was also apparent for other autoimmune disorders including narcolepsy, Crohn’s disease, systemic lupus erythematosus, and multiple sclerosis (MS) (**Fig. 5B** and **Supplemental Fig. S5A–C**), although the details were different for each trait. For instance, STAT1 in monocytes and STAT5B in T cells were commonly enriched for IBD and MS, whereas SPI1 in macrophages was specific for IBD and MYB in T cell–derived Jurkat cells was unique to MS. Given that SPI1 and MYB were enriched at macrophage- and T cell–specific enhancers, respectively (**Fig. 3E**,**F**), these data are consistent with the fact that inflammation in IBD and MS is largely mediated by intestinal macrophages and lymphocytes in the central nervous system, respectively (Fletcher et al. 2010; Lehmann-Horn et al. 2013; Steinbach and Plevy 2014). Although enrichment analysis with TF motifs or ChIP-seq data of ENCODE and Roadmap Epigenomics also showed immune-related TF enrichment for such diseases (Thurman et al. 2012; Schaub et al. 2012; Farh et al. 2014), our results reveal a greater variety of TFs such as SPI1 for IBD and MYB for MS, again highlighting the detection power conferred by the large amount of real binding data accessed by ChIP-Atlas.

**Figure 5.**
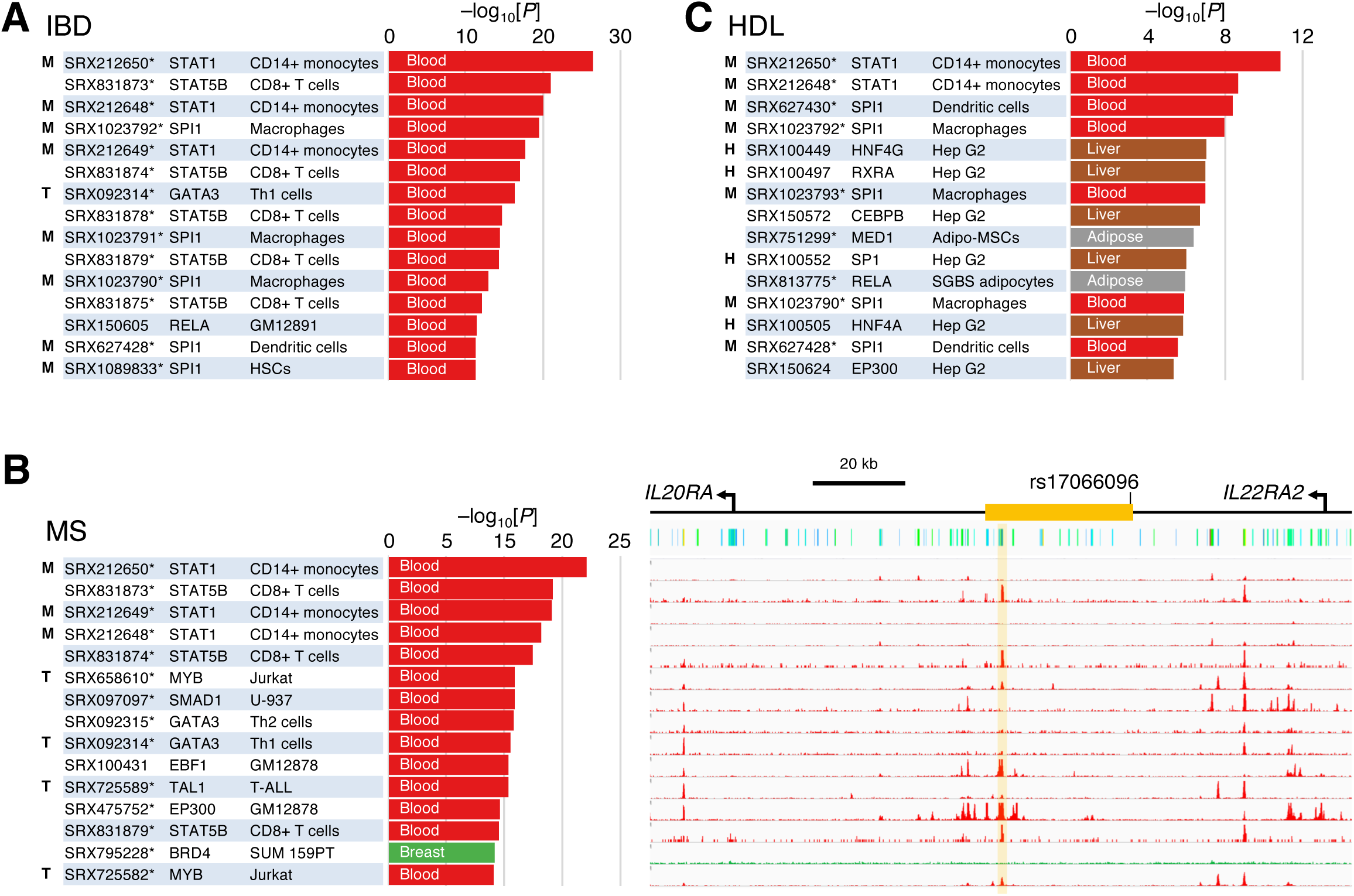
TFs enriched for both tissue-specific enhancers and noncoding GWAS-identified SNPs. (*A*–*C*) ChIP-seq data enriched for the regions associated with IBD (*A*), MS (*B*), and HDL cholesterol level (*C*) are shown as in **Fig. 4B**. Bold letters indicate the SRXs also enriched at enhancers specifically activated in hepatocytes (“H”), macrophages (“M”), and T cells (“T”) as shown in **Fig. 3C**,**E**,**F**, respectively.

It is unlikely that enrichment of breast- and blood-related TFs in the above results was due to corresponding larger amounts of ChIP-seq data (**Fig. 1C**), as indicated by our findings for high density lipoprotein (HDL)–cholesterol levels (**Fig. 5C**). In this instance, significantly enriched ChIP-seq data originated from the liver, adipose, and blood-related cells. The enriched TFs were similar to those enriched at hepatocyte and macrophage-specific enhancers (HNF4 and CEBPB in Hep G2 cells and SPI1 in macrophages, respectively; **Figs. 5C** and **3C**,**E**). These results are consistent with the cholesterol cycle, in which cholesterol is synthesized in the liver, stored in adipose tissue, and transported to peripheral tissues and cells such as macrophages, from which excess amounts are retrieved by HDL to protect against atherosclerosis (Tabas 2010; Yamamoto et al. 2016). Taking full advantage of ChIP-seq data in SRA thus supports the detection not only of TFs but also of cell types that contribute to human traits associated with noncoding variants. Similar to our results for BRC, we detected TF hotspots for other traits (**Fig. 5B** right; **Supplemental Fig. S6**), which may prove informative for the identification of causal variants and shed light on pathological mechanisms underlying the action of noncoding polymorphisms.

## DISCUSSION

Previous studies have attempted integrative functional annotation of enhancers and GWAS-identified SNPs in noncoding genomic regions with the use of ChIP-seq data for TFs from the ENCODE project (Gerstein et al. 2012; Schaub et al. 2012) or of data for active histone marks (Trynka et al. 2013; Farh et al. 2014) or TF motifs at DHSs (Maurano et al. 2012). We have now achieved this goal by making full use of chromatin-profiling data from SRA. We identified a large number of TFs enriched at tissue-specific enhancers, which are thought to serve as master regulators of corresponding cell types and are potential targets for cell reprograming. These results also provided insight into our GWAS analyses as exemplified by the finding that TFs enriched at blood-related enhancers were reproducibly enriched at LD-DHSs associated with autoimmune disorders, whereas those enriched at hepatocyte or macrophage enhancers also accumulated at LD-DHSs associated with HDL-cholesterol levels. Our two main types of analyses—TF-based functional annotation of enhancers and of SNPs identified by GWASs—were thus found to provide mutual support with regard to their physiological relevance. Of note, we identified hotspot regions containing LD-DHSs that were occupied by multiple TFs. Such a feature is a hallmark of enhancers with essential and robust activities (Hnisz et al. 2013), suggesting that the hotspots are functional regions in pathological processes. Fine-mapping of causal variants within the hotspot regions and evaluation of the physiological functions of the hotspots by genomic editing are thus warranted. In addition to the data presented above, we obtained plausible results for other GWAS analyses, including enrichment of cell types of the Prostate class for bladder cancer (both the prostate and bladder are derived from the urogenital sinus; **Supplemental Fig. S5D**), those of the Blood class for hematocyte-related traits (such as red blood cell traits, mean corpuscular hemoglobin or volume, and chronic lymphocytic leukemia), and of hormonal receptor–type TFs (ESR1 or AR) for traits related to hormone balance (such as temperament in bipolar disorder, height-adjusted body mass index, and migraine with aura) (see URL in **Supplemental Table S1** for the latest complete results). These data should prove to be informative with regard to the TFs and cell types directly engaged in the corresponding pathological mechanisms and processes.

Compared with other databases that have been developed to visualize individual ChIP-seq data (Sun et al. 2013; Yevshin et al. 2017; Yang et al. 2013), ChIP-Atlas not only contains a greater amount of data but also is characterized by the distinctive features that the data are fully integrated and processed for data-mining.

Regulatory factors for given genes and loci can thus be identified with the “Peak Browser” function, potential target genes of given TFs with “Target Genes,” and TF-TF interactions with “Colocalization” (**Fig. 1E**), all of which are fundamental to an understanding of gene regulatory networks. The fourth function of ChIP-Atlas, “in silico ChIP,” is useful to identify TFs enriched at given genomic ROIs, including regions associated with human traits and enhancers activated in specific cell types, as shown above, as well as other ROIs such as expression quantitative trait loci (eQTL), the user’s own ChIP-seq peak-call data, and evolutionally accelerated regions (e.g. Anan et al. 2018; Ferris et al. 2018; Ishigaki et al. 2017). This tool is also able to accept a batch of gene symbols and can therefore discover TFs that collectively regulate the gene set of interest (e.g. Chatterjee et al. 2018; Onodera et al. 2017; Chen et al. 2017). The results generated by ChIP-Atlas are all assigned unique URLs (**Supplemental Table S1**) and are publicly available, and they are thus ready for sharing seamlessly among researchers, for subsequent analysis in command-lines, and for interconnecting with other biodatabases such as DeepBlue epigenomic data server (Albrecht et al. 2016), where ChIP-Atlas data can be imported for analyses. Given that we designed ChIP-Atlas to be updated monthly with semiautomatic pipelines, the source data will be continuously expanded and may include more sophisticated chromatin-profiling results.

## MATERIALS AND METHODS

### Source data and primary processing

Sample metadata and biosample data of all SRXs were downloaded from NCBI FTP sites (ftp://ftp.ncbi.nlm.nih.gov/sra/reports/Metadata and ftp://ftp.ncbi.nlm.nih.gov/biosample). ChIP-Atlas uses SRXs that meet the following criteria: LIBRARY_STRATEGY is “ChIP-seq” or “DNase-Hypersensitivity”; LIBRARY_SOURCE is “GENOMIC”; taxonomy_name is “Homo sapiens,” “Mus musculus,” “Caenorhabditis elegans,” “Drosophila melanogaster,” or “Saccharomyces cerevisiae”; and INSTRUMENT_MODEL includes “Illumina” (the proportion of non-Illumina platforms is ~5%). Binarized sequence raw data (.sra) for each SRX were downloaded and decompressed into Fastq format with the “fastq-dump” command of SRA Toolkit (http://www.ncbi.nlm.nih.gov, ver. 2.3.2-4) according to a default mode, with the exception of paired-end reads, which were decoded with the “--split-files” option. In the case of an SRX including multiple runs, decompressed Fastq files were concatenated into a single such file. Fastq files were then aligned with the use of Bowtie2 (ver. 2.2.2) (Langmead and Salzberg 2012) according to a default mode, with the exception of paired-end reads, for which two Fastq files were specified with “-1” and “-2” options. The following genome assemblies were used for alignment and subsequent processing: hg19 (*H. sapiens*), mm9 (*M. musculus*), rn6 (*R*. *norvegicus*), dm3 (*D. melanogaster*), ce10 (*C. elegans*), and sacCer3 (*S. cerevisiae*). Resultant SAM-formatted files were binarized into BAM format with SAMtools (ver. 0.1.19; samtools view) (Li et al. 2009) and sorted (samtools sort) before removal of polymerase chain reaction (PCR) duplicates (samtools rmdup). BedGraph-formatted coverage scores were calculated with BEDTools (ver. 2.17.0; genomeCoverageBed) (Quinlan and Hall 2010) in RPM (reads per million mapped reads) units with the “-scale 1000000/N” option, where N is mapped read counts after removal of PCR duplicates. BedGraph files were binarized into BigWig format with the UCSC BedGraphToBigWig tool (http://hgdownload.cse.ucsc.edu, ver. 4). BAM files were also used for peak-calling with MACS2 (ver. 2.1.0, macs2 callpeak) (Zhang et al. 2008) in BED4 format. Options for *Q* value threshold (<1 × 10^−5^, <1 × 10^−10^, or <1 × 10^−20^) were applied, and the following options were set for genome size: “-g hs” (*H. sapiens*), “-g mm” (*M. musculus*), “-g 2.15e9” (*R*. *norvegicus*), “-g dm” (*D. melanogaster*), “-g ce” (*C. elegans*), and “-g 12100000” (*S. cerevisiae*).

### Curation of sample metadata

Sample metadata of all SRXs (biosample_set.xml) were downloaded (ftp://ftp.ncbi.nlm.nih.gov/biosample) to extract the attributes for antigens and antibodies (http://dbarchive.biosciencedbc.jp/kyushu-u/metadata/ag_attributes.txt) as well as cell types and tissues (http://dbarchive.biosciencedbc.jp/kyushu-u/metadata/ct_attributes.txt). For sorting out of metadata dependent on the discretion of the submitter, antigens and cell types were manually annotated by curators who had been fully trained in molecular and developmental biology. Each annotation has a “Class” and “Subclass” as shown in antigenList.tab and celltypeList.tab, which are available from the document page of ChIP-Atlas (https://github.com/inutano/chip-atlas/wiki). Guidelines for antigen annotation included the following: (1) histones were based on Brno nomenclature (such as H3K4me3 and H3K27ac) (Turner 2005); (2) gene symbols were used for gene-encoded human proteins according to HGNC (http://www.genenames.org) (for example, OCT3/4 → POU5F1, and p53 → TP53); and (3) modifications such as phosphorylation were ignored. If an antibody recognizes multiple molecules in a family, the first in an ascending order was chosen (for example, anti-SMAD2/3 → SMAD2).

Most human, mouse, and rat cell types were classified according to their tissue of origin. Embryonic stem cells and induced pluripotent stem cells were classified in the “Pluripotent stem cell” class. The nomenclature of cell lines and tissue names was standardized according to the following frameworks and resources: (1) Supplementary Table S2 of a paper (Yu et al. 2015) proposing unified cell line names (for example, MDA-231, MDA231, or MDAMB231 → MDA-MB-231); (2) ATCC, a nonprofit repository of cell lines (http://www.atcc.org); (3) cell line nomenclature used in the ENCODE project; and (4) MESH for tissue names (http://www.ncbi.nlm.nih.gov/mesh).

Antigens or cell types were categorized as “Unclassified” if the curators could not understand attribute values, and as “No description” if there was no attribute value, with these two classes being combined and designated “Unclassified” for simplicity in the present study (**Fig. 1B**,**C**). More details and examples are available from the document page of ChIP-Atlas (https://github.com/inutano/chip-atlas/wiki).

### Peak Browser

BED4-formatted peak-call data of each SRX were concatenated and converted to BED9 + GFF3 format to show MACS2 scores and sample metadata in the genome browser IGV. The descriptions are provided on our document page (https://github.com/inutano/chip-atlas/wiki).

### in silico ChIP

On submission, the two sets of BED-formatted data (ROIs and background regions, see below) are sent to the NIG supercomputer server, where the overlaps between the submitted data and peak-call data archived in ChIP-Atlas are counted with the “intersect” command of BEDTools2 (ver. 2.23.0). *P* values are calculated with the two-tailed Fisher’s exact probability test (the null hypothesis is that the two data sets overlap with the ChIP-Atlas peak-call data in the same proportion). *Q* values are calculated with the Benjamini and Hochberg method.

### Analysis of GWAS data

GWAS data were downloaded from the UCSC FTP site (http://hgdownload.cse.ucsc.edu/goldenPath/hg19/database/gwasCatalog.txt.gz), and LD-block data (phase 3 of the 1000 Genomes Project, European, *r*^2^ > 0.9) were from SNPsnap (http://www.broadinstitute.org/mpg/snpsnap/database/EUR/ld0.9/ld0.9_collection.tab.gz). LD-DHSs were prepared as described in **Supplemental Fig. S3A** and classified according to traits. They were submitted to the “*in silico* ChIP” Web page with the following parameter settings: “Antigen Class” is “TFs and others”; “Cell type Class” is “All cell types”; and “Threshold for Significance” (MACS2 score) is “100.” The background data for comparison were prepared by assembling LD-DHSs of all traits with the exception of the target trait.

### Analysis of tissue-specific enhancers

FANTOM5 facet-specific predefined enhancer data were downloaded from PrESSTo (http://pressto.binf.ku.dk) and submitted to the “*in silico* ChIP” Web page with the following parameter settings: “Antigen Class” is “TFs and others”; “Cell type Class” is “All cell types”; and “Threshold for Significance” (MACS2 score) is “100.” The background data for comparison were prepared by assembling FANTOM5 enhancers of all facets with the exception of the target facet.

### Cluster analysis

Each matrix of LD-DHSs and SRXs was binarized to 1 (overlapped) or 0 (nonoverlapped), and cluster analysis was performed with the R package (dist, method = “euclidean”; hclust, method = “complete”).

## ACKNOWLEDGMENTS

The public data used in this research were obtained from SRA of NCBI. Computations were performed mostly on the NIG supercomputer at ROIS National Institute of Genetics. This work was supported by KAKENHI grants 25840087, 26291051, 17H01571, and 15K14529 from the Japan Society for the Promotion of Science (JSPS), and by the National Bioscience Database Center (NBDC) of the Japan Science and Technology Agency (JST).

