## Supplementary Materials for "Integrative analysis of transcription factor occupancy at enhancers and disease risk loci in noncoding genomic regions" (from another version of this preprint: version 2, 2018-03-31)

**Supplementary Figure 1.** Sample and library metadata curated in ChIP-Atlas. (*A*) Sample metadata for all ChIP-seq and DNase-seq data in ChIP-Atlas are systematically curated, allowing the data to be searched with given key words such as antigen and cell type names. (*B*) Snapshot of the Web page for ChIP-Atlas "Peak Browser" function is shown, with the results of the settings being presented in **Fig. 2**. (*C*) Example of the detailed information for SRX187209, including the sample metadata described by ChIP-Atlas curators and the original data submitter, processing logs, and read quality from DBCLS SRA (http://sra.dbcls.jp). Blue buttons at the top are controllers for showing the alignment and peak-call data in IGV ("View on IGV"), for downloading these data ("Download"), for viewing the analyzed data by ChIP-Atlas "Target Genes" and "Colocalization" ("View Analysis"), and for opening the external pages showing details for the experimental conditions and materials ("Link Out"). This type of Web page appears on clicking the bars in the "Peak Browser" view (**Fig. 2**) as well as by clicking SRX IDs shown in Web pages for a key word search (*A*) or "in silico ChIP" results (**Supplementary Fig. S2**).

**Supplementary Figure 2.** Example of analysis by "*in silico* ChIP." At the Web site of "*in silico* ChIP" (*A*), a user can submit two sets of genomic intervals in BED format: data of interest in the orange area and background data for comparison in the gray area. It is also possible to filter the results according to antigen and cell type classes as well as to set a threshold for the MACS2 score. On clicking the "submit" button, the data are sent to an NIG supercomputer server for performance of the enrichment analysis, as shown in **Fig. 3A**. In several minutes, a table is returned to the user's terminal as exemplified in (*B*), which shows the results for submission in (*A*) of "TFs and others" for "1. Antigen Class"; "All cell types" for "2. Cell type Class"; "100" for "3. Threshold for Significance"; FANTOM5 hepatocyte-specific enhancers for "4. Select your data"; and FANTOM5 tissue-specific enhancers except hepatocyte enhancers for "5. Select dataset to be compared." The first row in (*B*), for example, indicates EP300 ChIP-seq data (SRX100544) for Hep G2 cells. The total number of peaks for EP300 is 24,334, of which 80 peaks overlap with hepatocyte-specific enhancers ($n$ = 286) and 1,147 peaks overlap with other enhancers ($n$ = 20,509), yielding a $P$ value of $1 \times 10^{-32.1}$ (Fisher's exact probability test), $Q$ value of $1 \times 10^{-28.4}$ (Benjamini & Hochberg method), and fold enrichment of 5.00. The table is sorted according to $P$ value, where HNF4A/G and

FOXA1/2 in Hep G2 are ranked at 4th, 8th, 11th, and 13th. The table is also graphically summarized in **Fig. 3C**, in which each row of the table is represented by a bar to indicate the *P* value.

**Supplementary Figure 3.** Characteristics of LD-DHSs. (*A*) Concept and delineation of LD-DHSs. (1) A BED file of the marker SNP loci (X marks) associated with GWAS traits was obtained from the NHGRI-EBI GWAS catalog. (2) The LD blocks (medium blue) (phase 3 of the 1000 Genomes Project, European, $r^2 > 0.9$) corresponding to each SNP locus were extended by 1 kb at each end (pale blue). (3) Genomic features in (2) were removed if they overlapped with exons (gray) of RefSeq coding or noncoding genes. (4) Regions of overlap between the remaining features in (3) and all DHSs in all cell types recorded in ChIP-Atlas (green, MACS2 score > 100) were extracted and used as LD-DHSs (orange). (*B*) Aggregation plots (bin size of 100 bp) for the average number of peaks for human TFs recorded in ChIP-Atlas (MACS2 score > 100) that are located around loci for GWAS marker SNPs and LD-DHSs (blue and orange, respectively). Randomly permutated data are shown as the background ("Shuffled," gray). (*C*) Histogram showing the distribution of the size of LD blocks (phase 3 of the 1000 Genomes Project, European, $r^2 > 0.9$) for GWAS marker SNPs (blue) and LD-DHSs (orange). Note that TF binding was rarely enriched at GWAS marker SNPs themselves (*B*), likely reflecting the underlying GWAS hypothesis that most causal variants do not coincide with marker SNP loci but rather lie within the corresponding LD blocks. Given that most LD blocks range in size from tens to hundreds of kilobases (*C*), large portions of these blocks would not be expected to be related to either causal variants or regulatory functions. To focus on the regulatory elements in these noncoding regions, we therefore introduced the concept of LD-DHSs, defined as DHSs located within noncoding regions that are in LD with GWAS SNPs (*A*). LD-DHSs are only hundreds of bases in length (*C*) and are enriched in multiple TFs (*B*).

**Supplementary Figure 4.** LD-DHS hotspots where multiple TFs accumulate. LD-DHSs for breast cancer (BRC) and the significantly enriched SRXs ($Q < 0.01$) (corresponding to **Fig. 4B**) are shown in columns and rows, respectively. When an LD-DHS overlaps with the ChIP-seq peaks of an SRX, the corresponding cell type in the matrix is colored according to the cell type class shown in **Fig. 1C**, with rows and columns of the matrix clustered hierarchically. The top 14 SRXs and LD-DHSs shown in **Fig. 4B** are labeled on the left and bottom, respectively.

**Supplementary Figure 5.** TFs enriched at noncoding GWAS SNPs. Results for TF enrichment analysis with "*in silico* ChIP" for narcolepsy (*A*), systemic lupus erythematosus (SLE) (*B*), Crohn's disease (CD) (*C*), and bladder cancer (BLC) (*D*) are shown.

**Supplementary Figure 6.** Examples of hotspots around noncoding GWAS SNPs. Alignment data for the ChIP-seq experiments indicated on the left and for the regions around marker SNPs associated with IBD (*A*) and HDL-cholesterol levels (*C*) are shown, with the *y*-axes ranging from 0 to 1 RPM units.

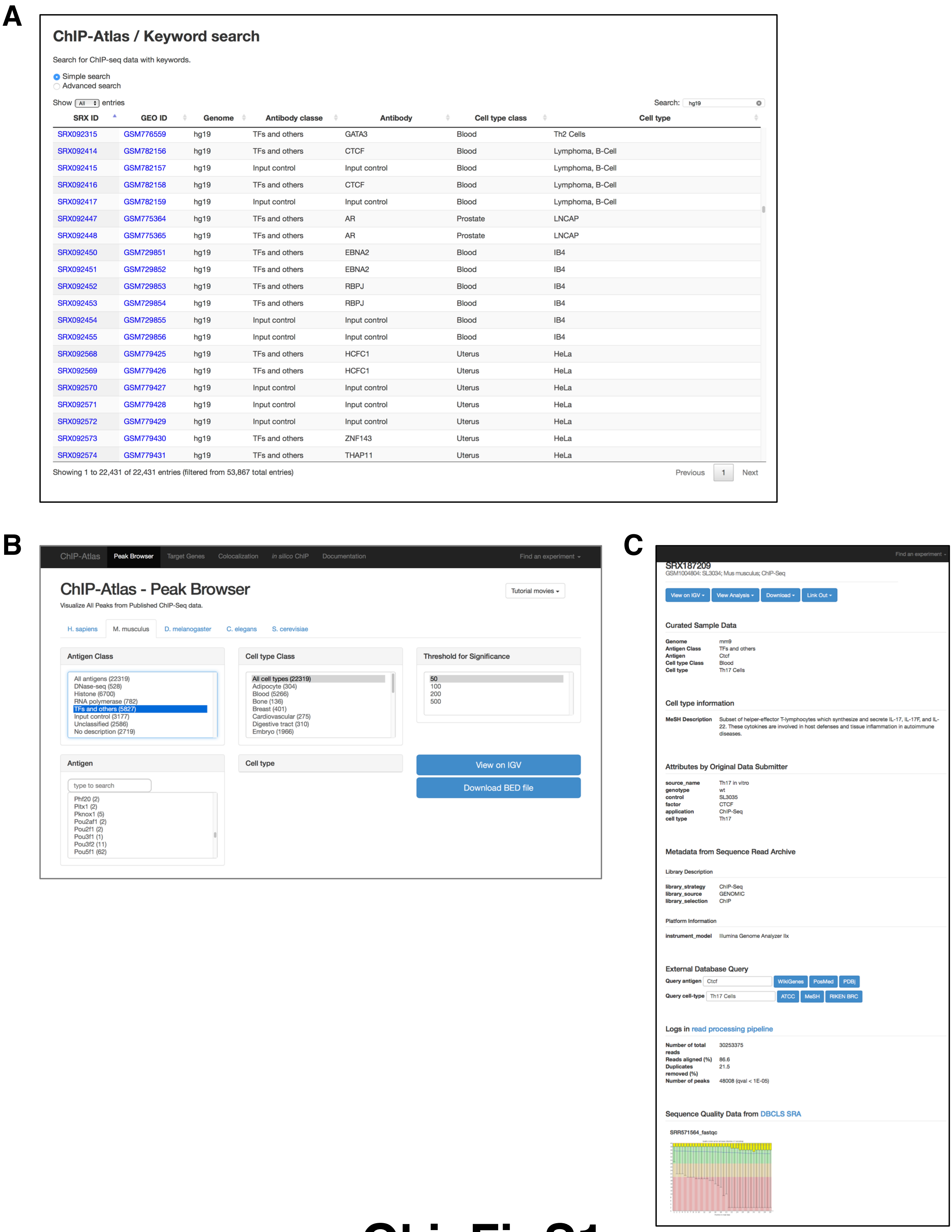
A
ChIP-Atlas / Keyword search
Search for ChIP-seq data with keywords.
Simple search
Advanced search
Show All entries
Search: hg19
SRX ID | GEO ID | Genome | Antibody classe | Antibody | Cell type class | Cell type
SRX092315 GSM776559 hg19 TFs and others GATA3 Blood Th2 Cells
SRX092414 GSM782156 hg19 TFs and others CTCF Blood Lymphoma, B-Cell
SRX092415 GSM782157 hg19 Input control Input control Blood Lymphoma, B-Cell
SRX092416 GSM782158 hg19 TFs and others CTCF Blood Lymphoma, B-Cell
SRX092417 GSM782159 hg19 Input control Input control Blood Lymphoma, B-Cell
SRX092447 GSM775364 hg19 TFs and others AR Prostate LNCAP
SRX092448 GSM775365 hg19 TFs and others AR Prostate LNCAP
SRX092450 GSM729851 hg19 TFs and others EBNA2 Blood IB4
SRX092451 GSM729852 hg19 TFs and others EBNA2 Blood IB4
SRX092452 GSM729853 hg19 TFs and others RBPJ Blood IB4
SRX092453 GSM729854 hg19 TFs and others RBPJ Blood IB4
SRX092454 GSM729855 hg19 Input control Input control Blood IB4
SRX092455 GSM729856 hg19 Input control Input control Blood IB4
SRX092568 GSM779425 hg19 TFs and others HCFC1 Uterus HeLa
SRX092569 GSM779426 hg19 TFs and others HCFC1 Uterus HeLa
SRX092570 GSM779427 hg19 Input control Input control Uterus HeLa
SRX092571 GSM779428 hg19 Input control Input control Uterus HeLa
SRX092572 GSM779429 hg19 Input control Input control Uterus HeLa
SRX092573 GSM779430 hg19 TFs and others ZNF143 Uterus HeLa
SRX092574 GSM779431 hg19 TFs and others THAP11 Uterus HeLa
Showing 1 to 22,431 of 22,431 entries (filtered from 53,867 total entries)
Previous 1 Next
B
ChIP-Atlas Peak Browser Target Genes Colocalization in silico ChIP Documentation Find an experiment
ChIP-Atlas - Peak Browser
Tutorial movies
Visualize All Peaks from Published ChIP-Seq data.
H. sapiens M. musculus D. melanogaster C. elegans S. cerevisiae
Antigen Class
All antigens (22319)
DNase-seq (528)
Histone (6700)
RNA polymerase (782)
TFs and others (5827)
Input control (3177)
Unclassified (2586)
No description (2719)
Cell type Class
All cell types (22319)
Adipocyte (304)
Blood (5266)
Bone (136)
Breast (401)
Cardiovascular (275)
Digestive tract (310)
Embryo (1966)
Threshold for Significance
50
100
200
500
Antigen
type to search
Phf20 (2)
Pitx1 (2)
Pknox1 (5)
Pou2af1 (2)
Pou2f1 (2)
Pou3f1 (1)
Pou3f2 (11)
Pou5f1 (62)
Cell type
View on IGV
Download BED file
C
Find an experiment
SRX187209
GSM1004804: SL3034; Mus musculus; ChIP-Seq
View on IGV View Analysis Download Link Out
Curated Sample Data
Genome mm9
Antigen Class TFs and others
Antigen Ctcf
Cell type Class Blood
Cell type Th17 Cells
Cell type information
MeSH Description Subset of helper-effector T-lymphocytes which synthesize and secrete IL-17, IL-17F, and IL-22. These cytokines are involved in host defenses and tissue inflammation in autoimmune diseases.
Attributes by Original Data Submitter
source_name Th17 in vitro
genotype wt
control SL3035
factor CTCF
application ChIP-Seq
cell type Th17
Metadata from Sequence Read Archive
Library Description
library_strategy ChIP-Seq
library_source GENOMIC
library_selection ChIP
Platform Information
instrument_model Illumina Genome Analyzer IIx
External Database Query
Query antigen Ctcf WikiGenes PosMed PDBj
Query cell-type Th17 Cells ATCC MeSH RIKEN BRC
Logs in read processing pipeline
Number of total reads 30253375
Reads aligned (%) 86.6
Duplicates removed (%) 21.5
Number of peaks 48008 (qval < 1E-05)
Sequence Quality Data from DBCLS SRA
SRR571564_fastqc

**Oki_FigS1**

A

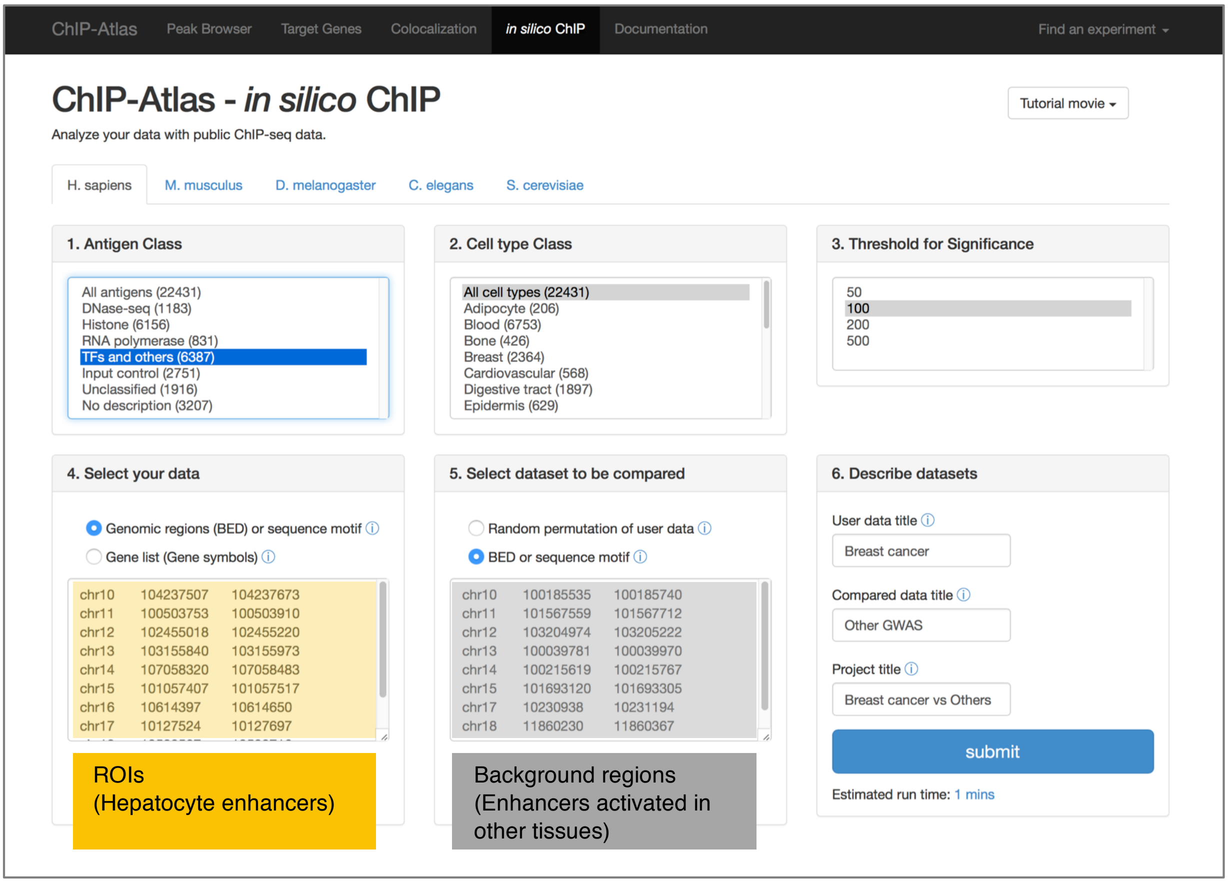


B

| ID | Antigen class | Antigen | Cell class | Cell | Num of peaks | Overlaps / Hepatocyte | Overlaps / Other enhancers | Log P-val | Log Q-val | Fold Enrichment | FE > 1? |
|---|---|---|---|---|---|---|---|---|---|---|---|
| SRX100544 | TFs and others | EP300 | Liver | Hep G2 | 24334 | 80/286 | 1147/20509 | -32.1 | -28.4 | 5.00 | TRUE |
| SRX190321 | TFs and others | MAX | Liver | Hep G2 | 40220 | 90/286 | 1497/20509 | -31.9 | -28.4 | 4.31 | TRUE |
| SRX100552 | TFs and others | SP1 | Liver | Hep G2 | 19032 | 64/286 | 755/20509 | -29.7 | -26.4 | 6.08 | TRUE |
| SRX100449 | TFs and others | HNF4G | Liver | Hep G2 | 15919 | 54/286 | 507/20509 | -29.3 | -26.2 | 7.64 | TRUE |
| SRX100497 | TFs and others | RXRA | Liver | Hep G2 | 13022 | 54/286 | 557/20509 | -27.5 | -24.5 | 6.95 | TRUE |
| SRX100493 | TFs and others | HEY1 | Liver | Hep G2 | 26412 | 69/286 | 984/20509 | -27.5 | -24.5 | 5.03 | TRUE |
| SRX100538 | TFs and others | HDAC2 | Liver | Hep G2 | 16071 | 58/286 | 676/20509 | -27.0 | -24.1 | 6.15 | TRUE |
| SRX100505 | TFs and others | HNF4A | Liver | Hep G2 | 21259 | 54/286 | 585/20509 | -26.5 | -23.7 | 6.62 | TRUE |
| SRX190332 | TFs and others | MYBL2 | Liver | Hep G2 | 15213 | 55/286 | 637/20509 | -25.7 | -22.9 | 6.19 | TRUE |
| SRX190264 | TFs and others | CREB1 | Liver | Hep G2 | 26690 | 68/286 | 1092/20509 | -24.3 | -21.5 | 4.47 | TRUE |
| SRX100448 | TFs and others | FOXA2 | Liver | Hep G2 | 45130 | 67/286 | 1138/20509 | -22.6 | -19.9 | 4.22 | TRUE |
| SRX150360 | TFs and others | TBP | Liver | Hep G2 | 10293 | 38/286 | 327/20509 | -21.7 | -19.1 | 8.33 | TRUE |
| SRX100506 | TFs and others | FOXA1 | Liver | Hep G2 | 50941 | 70/286 | 1295/20509 | -21.7 | -19.0 | 3.88 | TRUE |

Showing 1 to 100 of 5,738 entries    Previous 1 2 3 4 5 … 58 Next

### Oki_FigS2

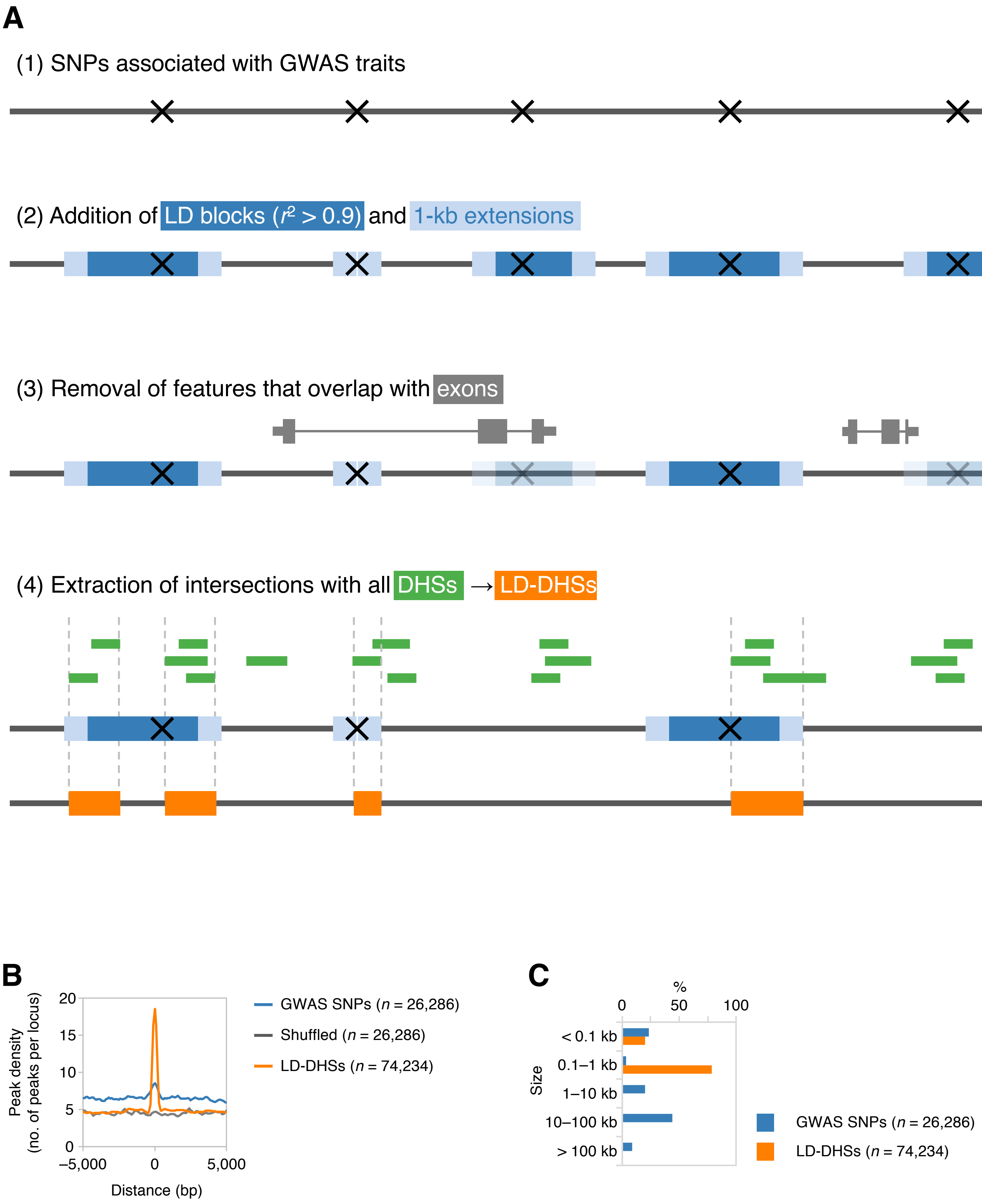
A
(1) SNPs associated with GWAS traits
(2) Addition of LD blocks ($r^2$ > 0.9) and 1-kb extensions
(3) Removal of features that overlap with exons
(4) Extraction of intersections with all DHSs → LD-DHSs
B
Peak density (no. of peaks per locus)
20
15
10
5
0
–5,000
0
5,000
Distance (bp)
GWAS SNPs ($n$ = 26,286)
Shuffled ($n$ = 26,286)
LD-DHSs ($n$ = 74,234)
C
%
0
50
100
Size
< 0.1 kb
0.1–1 kb
1–10 kb
10–100 kb
> 100 kb
GWAS SNPs ($n$ = 26,286)
LD-DHSs ($n$ = 74,234)

### Oki_FigS3

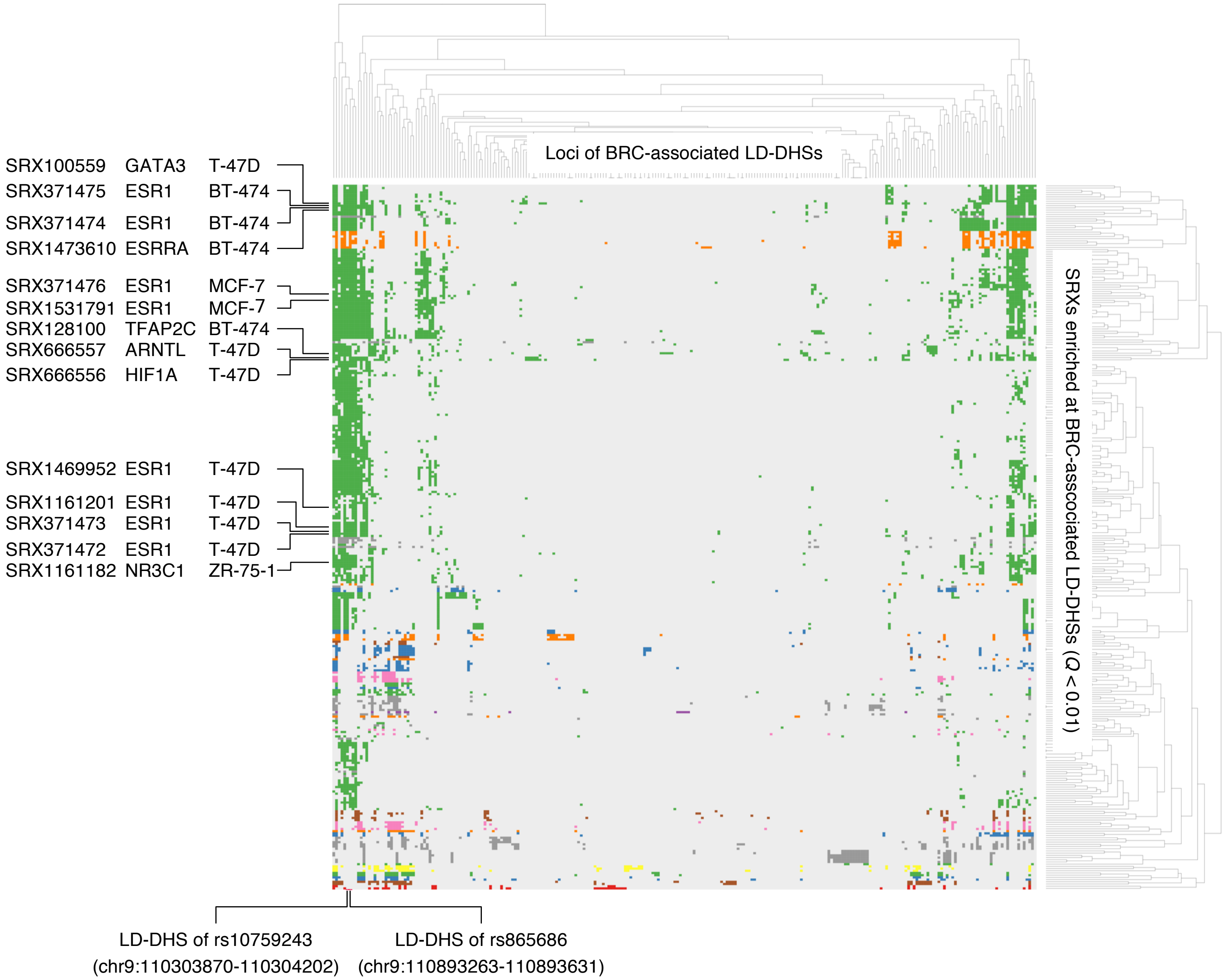
Loci of BRC-associated LD-DHSs
SRX100559 GATA3 T-47D
SRX371475 ESR1 BT-474
SRX371474 ESR1 BT-474
SRX1473610 ESRRA BT-474
SRX371476 ESR1 MCF-7
SRX1531791 ESR1 MCF-7
SRX128100 TFAP2C BT-474
SRX666557 ARNTL T-47D
SRX666556 HIF1A T-47D
SRX1469952 ESR1 T-47D
SRX1161201 ESR1 T-47D
SRX371473 ESR1 T-47D
SRX371472 ESR1 T-47D
SRX1161182 NR3C1 ZR-75-1
SRXs enriched at BRC-associated LD-DHSs (Q < 0.01)
LD-DHS of rs10759243
(chr9:110303870-110304202)
LD-DHS of rs865686
(chr9:110893263-110893631)

### Oki_FigS4

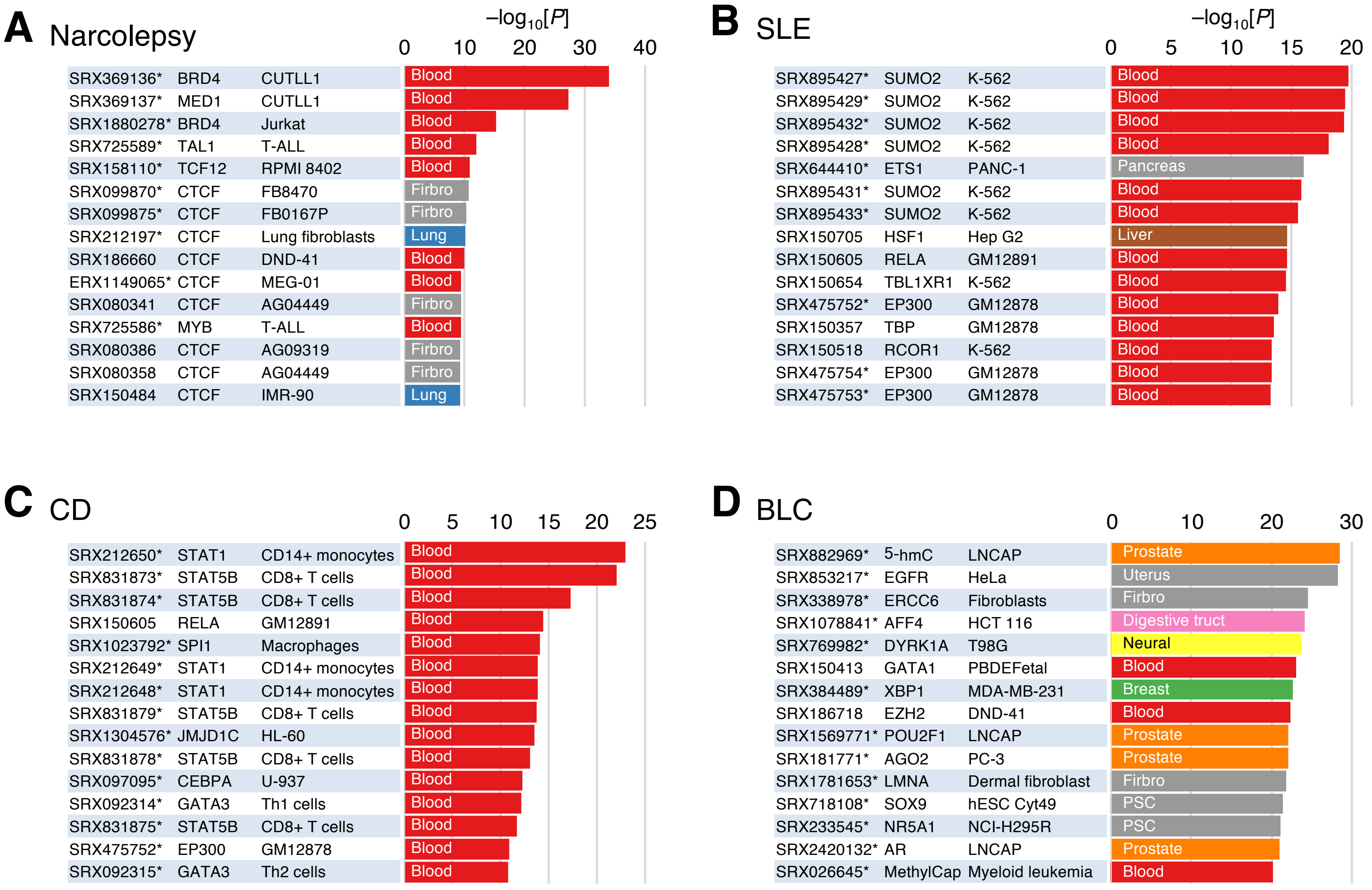

A Narcolepsy
–log10[P]
0 10 20 30 40
SRX369136* BRD4 CUTLL1 Blood
SRX369137* MED1 CUTLL1 Blood
SRX1880278* BRD4 Jurkat Blood
SRX725589* TAL1 T-ALL Blood
SRX158110* TCF12 RPMI 8402 Blood
SRX099870* CTCF FB8470 Firbro
SRX099875* CTCF FB0167P Firbro
SRX212197* CTCF Lung fibroblasts Lung
SRX186660 CTCF DND-41 Blood
ERX1149065* CTCF MEG-01 Blood
SRX080341 CTCF AG04449 Firbro
SRX725586* MYB T-ALL Blood
SRX080386 CTCF AG09319 Firbro
SRX080358 CTCF AG04449 Firbro
SRX150484 CTCF IMR-90 Lung
B SLE
–log10[P]
0 5 10 15 20
SRX895427* SUMO2 K-562 Blood
SRX895429* SUMO2 K-562 Blood
SRX895432* SUMO2 K-562 Blood
SRX895428* SUMO2 K-562 Blood
SRX644410* ETS1 PANC-1 Pancreas
SRX895431* SUMO2 K-562 Blood
SRX895433* SUMO2 K-562 Blood
SRX150705 HSF1 Hep G2 Liver
SRX150605 RELA GM12891 Blood
SRX150654 TBL1XR1 K-562 Blood
SRX475752* EP300 GM12878 Blood
SRX150357 TBP GM12878 Blood
SRX150518 RCOR1 K-562 Blood
SRX475754* EP300 GM12878 Blood
SRX475753* EP300 GM12878 Blood
C CD
0 5 10 15 20 25
SRX212650* STAT1 CD14+ monocytes Blood
SRX831873* STAT5B CD8+ T cells Blood
SRX831874* STAT5B CD8+ T cells Blood
SRX150605 RELA GM12891 Blood
SRX1023792* SPI1 Macrophages Blood
SRX212649* STAT1 CD14+ monocytes Blood
SRX212648* STAT1 CD14+ monocytes Blood
SRX831879* STAT5B CD8+ T cells Blood
SRX1304576* JMJD1C HL-60 Blood
SRX831878* STAT5B CD8+ T cells Blood
SRX097095* CEBPA U-937 Blood
SRX092314* GATA3 Th1 cells Blood
SRX831875* STAT5B CD8+ T cells Blood
SRX475752* EP300 GM12878 Blood
SRX092315* GATA3 Th2 cells Blood
D BLC
0 10 20 30
SRX882969* 5-hmC LNCAP Prostate
SRX853217* EGFR HeLa Uterus
SRX338978* ERCC6 Fibroblasts Firbro
SRX1078841* AFF4 HCT 116 Digestive truct
SRX769982* DYRK1A T98G Neural
SRX150413 GATA1 PBDEFetal Blood
SRX384489* XBP1 MDA-MB-231 Breast
SRX186718 EZH2 DND-41 Blood
SRX1569771* POU2F1 LNCAP Prostate
SRX181771* AGO2 PC-3 Prostate
SRX1781653* LMNA Dermal fibroblast Firbro
SRX718108* SOX9 hESC Cyt49 PSC
SRX233545* NR5A1 NCI-H295R PSC
SRX2420132* AR LNCAP Prostate
SRX026645* MethylCap Myeloid leukemia Blood


**Oki_FigS5**

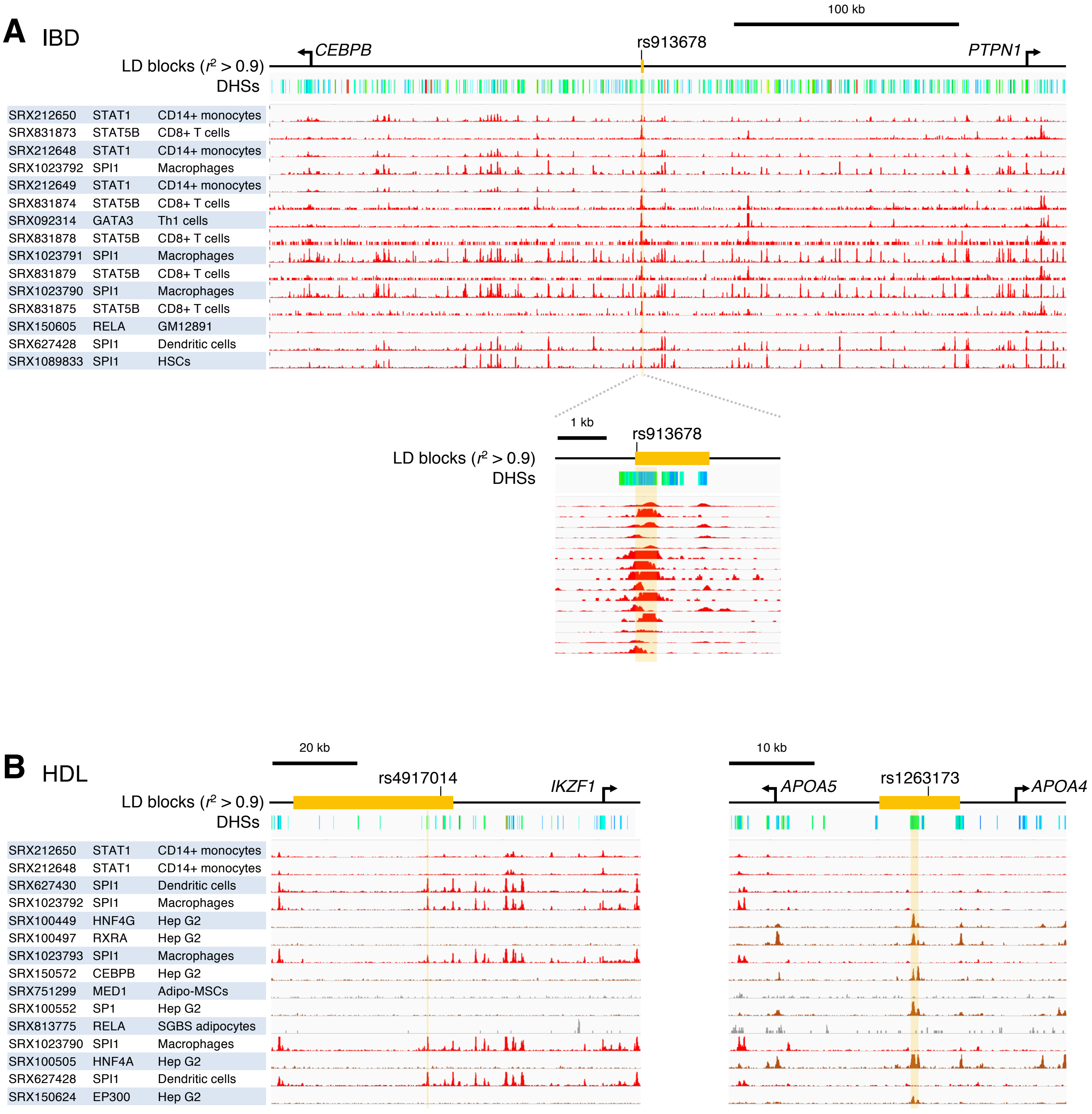
A
IBD
100 kb
CEBPB
rs913678
PTPN1
LD blocks (r2 > 0.9)
DHSs
SRX212650 STAT1 CD14+ monocytes
SRX831873 STAT5B CD8+ T cells
SRX212648 STAT1 CD14+ monocytes
SRX1023792 SPI1 Macrophages
SRX212649 STAT1 CD14+ monocytes
SRX831874 STAT5B CD8+ T cells
SRX092314 GATA3 Th1 cells
SRX831878 STAT5B CD8+ T cells
SRX1023791 SPI1 Macrophages
SRX831879 STAT5B CD8+ T cells
SRX1023790 SPI1 Macrophages
SRX831875 STAT5B CD8+ T cells
SRX150605 RELA GM12891
SRX627428 SPI1 Dendritic cells
SRX1089833 SPI1 HSCs
1 kb
rs913678
LD blocks (r2 > 0.9)
DHSs
B
HDL
20 kb
rs4917014
IKZF1
10 kb
APOA5
rs1263173
APOA4
LD blocks (r2 > 0.9)
DHSs
SRX212650 STAT1 CD14+ monocytes
SRX212648 STAT1 CD14+ monocytes
SRX627430 SPI1 Dendritic cells
SRX1023792 SPI1 Macrophages
SRX100449 HNF4G Hep G2
SRX100497 RXRA Hep G2
SRX1023793 SPI1 Macrophages
SRX150572 CEBPB Hep G2
SRX751299 MED1 Adipo-MSCs
SRX100552 SP1 Hep G2
SRX813775 RELA SGBS adipocytes
SRX1023790 SPI1 Macrophages
SRX100505 HNF4A Hep G2
SRX627428 SPI1 Dendritic cells
SRX150624 EP300 Hep G2

**Oki_FigS6**

| Figure | Description | URL |
|---|---|---|
| Figure 2 | Integrated peak-call data | http://dbarchive.biosciencedbc.jp/kyushu-u/mm9/assembled/Oth.ALL.10.AllAg.AllCell.bed |
| Figure 2 | SRX265314: Cebpb, Liver | http://dbarchive.biosciencedbc.jp/kyushu-u/mm9/eachData/bw/SRX265314.bw |
| Figure 2 | SRX187209: Ctcf, Th17 | http://dbarchive.biosciencedbc.jp/kyushu-u/mm9/eachData/bw/SRX187209.bw |
| Figure 3B | Complete data sets for enrichment analyses for enhancers | http://dbarchive.biosciencedbc.jp/kyushu-u/isc/FFenhancer.html |
| Figure 3C | Hepatocyte enhancers | http://dbarchive.biosciencedbc.jp/kyushu-u/isc/CL:0000182.html |
| Figure 3D | Blood vessel endothelial cell enhancers | http://dbarchive.biosciencedbc.jp/kyushu-u/isc/CL:0000071.html |
| Figure 3E | Macrophage enhancers | http://dbarchive.biosciencedbc.jp/kyushu-u/isc/CL:0000235.html |
| Figure 3F | T cell enhancers | http://dbarchive.biosciencedbc.jp/kyushu-u/isc/CL:0000084.html |
| Figure 4A | Complete data sets for enrichment analyses for GWAS | http://dbarchive.biosciencedbc.jp/kyushu-u/isc/gwas.html |
| Figure 4B | Enrichment analysis for BRC | http://dbarchive.biosciencedbc.jp/kyushu-u/isc/GWAS:0244.html |
| Figure 4B | SRX371474: ESR1, BT-474 | http://dbarchive.biosciencedbc.jp/kyushu-u/hg19/eachData/bw/SRX371474.bw |
| Figure 4B | SRX371475: ESR1, BT-474 | http://dbarchive.biosciencedbc.jp/kyushu-u/hg19/eachData/bw/SRX371475.bw |
| Figure 4B | SRX1469952: ESR1, T-47D | http://dbarchive.biosciencedbc.jp/kyushu-u/hg19/eachData/bw/SRX1469952.bw |
| Figure 4B | SRX666557: ARNTL, T-47D | http://dbarchive.biosciencedbc.jp/kyushu-u/hg19/eachData/bw/SRX666557.bw |
| Figure 4B | SRX1473610: ESRRA, BT-474 | http://dbarchive.biosciencedbc.jp/kyushu-u/hg19/eachData/bw/SRX1473610.bw |
| Figure 4B | SRX1531791: ESR1, MCF-7 | http://dbarchive.biosciencedbc.jp/kyushu-u/hg19/eachData/bw/SRX1531791.bw |
| Figure 4B | SRX371476: ESR1, MCF-7 | http://dbarchive.biosciencedbc.jp/kyushu-u/hg19/eachData/bw/SRX371476.bw |
| Figure 4B | SRX128100: TFAP2C, BT-474 | http://dbarchive.biosciencedbc.jp/kyushu-u/hg19/eachData/bw/SRX128100.bw |
| Figure 4B | SRX100559: GATA3, T-47D | http://dbarchive.biosciencedbc.jp/kyushu-u/hg19/eachData/bw/SRX100559.bw |
| Figure 4B | SRX666556: HIF1A, T-47D | http://dbarchive.biosciencedbc.jp/kyushu-u/hg19/eachData/bw/SRX666556.bw |
| Figure 4B | SRX1161182: NR3C1, ZR-75-1 | http://dbarchive.biosciencedbc.jp/kyushu-u/hg19/eachData/bw/SRX1161182.bw |
| Figure 4B | SRX371472: ESR1, T-47D | http://dbarchive.biosciencedbc.jp/kyushu-u/hg19/eachData/bw/SRX371472.bw |
| Figure 4B | SRX1161201: ESR1, T-47D | http://dbarchive.biosciencedbc.jp/kyushu-u/hg19/eachData/bw/SRX1161201.bw |
| Figure 4B | SRX371473: ESR1, T-47D | http://dbarchive.biosciencedbc.jp/kyushu-u/hg19/eachData/bw/SRX371473.bw |
| Figure 4B | SRX371447: ESR1, Tumour tissues | http://dbarchive.biosciencedbc.jp/kyushu-u/hg19/eachData/bw/SRX371447.bw |
| Figure 5A | Enrichment analysis for IBD | http://dbarchive.biosciencedbc.jp/kyushu-u/isc/GWAS:0663.html |
| Figure 5B | Enrichment analysis for MS | http://dbarchive.biosciencedbc.jp/kyushu-u/isc/GWAS:0817.html |
| Figure 5B | SRX212650: STAT1, CD14+ monocytes | http://dbarchive.biosciencedbc.jp/kyushu-u/hg19/eachData/bw/SRX212650.bw |
| Figure 5B | SRX831873: STAT5B, CD8+ T cells | http://dbarchive.biosciencedbc.jp/kyushu-u/hg19/eachData/bw/SRX831873.bw |
| Figure 5B | SRX212649: STAT1, CD14+ monocytes | http://dbarchive.biosciencedbc.jp/kyushu-u/hg19/eachData/bw/SRX212649.bw |
| Figure 5B | SRX212648: STAT1, CD14+ monocytes | http://dbarchive.biosciencedbc.jp/kyushu-u/hg19/eachData/bw/SRX212648.bw |
| Figure 5B | SRX831874: STAT5B, CD8+ T cells | http://dbarchive.biosciencedbc.jp/kyushu-u/hg19/eachData/bw/SRX831874.bw |
| Figure 5B | SRX658610: MYB, Jurkat | http://dbarchive.biosciencedbc.jp/kyushu-u/hg19/eachData/bw/SRX658610.bw |
| Figure 5B | SRX097097: SMAD1, U-937 | http://dbarchive.biosciencedbc.jp/kyushu-u/hg19/eachData/bw/SRX097097.bw |
| Figure 5B | SRX092315: GATA3, Th2 cells | http://dbarchive.biosciencedbc.jp/kyushu-u/hg19/eachData/bw/SRX092315.bw |
| Figure 5B | SRX092314: GATA3, Th1 cells | http://dbarchive.biosciencedbc.jp/kyushu-u/hg19/eachData/bw/SRX092314.bw |
| Figure 5B | SRX100431: EBF1, GM12878 | http://dbarchive.biosciencedbc.jp/kyushu-u/hg19/eachData/bw/SRX100431.bw |
| Figure 5B | SRX725589: TAL1, T-ALL | http://dbarchive.biosciencedbc.jp/kyushu-u/hg19/eachData/bw/SRX725589.bw |
| Figure 5B | SRX475752: EP300, GM12878 | http://dbarchive.biosciencedbc.jp/kyushu-u/hg19/eachData/bw/SRX475752.bw |
| Figure 5B | SRX831879: STAT5B, CD8+ T cells | http://dbarchive.biosciencedbc.jp/kyushu-u/hg19/eachData/bw/SRX831879.bw |
| Figure 5B | SRX795228: BRD4, SUM 159PT | http://dbarchive.biosciencedbc.jp/kyushu-u/hg19/eachData/bw/SRX795228.bw |
| Figure 5B | SRX725582: MYB, Jurkat | http://dbarchive.biosciencedbc.jp/kyushu-u/hg19/eachData/bw/SRX725582.bw |
| Figure 5C | Enrichment analysis for HDL | http://dbarchive.biosciencedbc.jp/kyushu-u/isc/GWAS:0563.html |
| Figure S1A | ChIP-Atlas, keyword search | http://dbarchive.biosciencedbc.jp/kyushu-u/metadata/refineSearchList.html |
| Figure S1B | ChIP-Atlas, Peak Browser | http://chip-atlas.org/peak_browser |
| Figure S1C | Details for SRX187209 | http://chip-atlas.org/view?id=SRX187209 |
| Figure S2A | ChIP-Atlas, in silico ChIP | http://chip-atlas.org/in_silico_chip |
| Figure S2B | Enrichment analysis for hepatocyte enhancers | http://dbarchive.biosciencedbc.jp/kyushu-u/isc/CL:0000182.html |
| Figure S5A | Enrichment analysis for Narcolepsy | http://dbarchive.biosciencedbc.jp/kyushu-u/isc/GWAS:0832.html |
| Figure S5B | Enrichment analysis for SLE | http://dbarchive.biosciencedbc.jp/kyushu-u/isc/GWAS:1239.html |
| Figure S5C | Enrichment analysis for CD | http://dbarchive.biosciencedbc.jp/kyushu-u/isc/GWAS:0376.html |
| Figure S5D | Enrichment analysis for BLC | http://dbarchive.biosciencedbc.jp/kyushu-u/isc/GWAS:0195.html |
| Figure S6A | SRX212650: STAT1, CD14+ monocytes | http://dbarchive.biosciencedbc.jp/kyushu-u/hg19/eachData/bw/SRX212650.bw |
| Figure S6A | SRX831873: STAT5B, CD8+ T cells | http://dbarchive.biosciencedbc.jp/kyushu-u/hg19/eachData/bw/SRX831873.bw |
| Figure S6A | SRX212648: STAT1, CD14+ monocytes | http://dbarchive.biosciencedbc.jp/kyushu-u/hg19/eachData/bw/SRX212648.bw |
| Figure S6A | SRX1023792: SPI1, Macrophages | http://dbarchive.biosciencedbc.jp/kyushu-u/hg19/eachData/bw/SRX1023792.bw |
| Figure S6A | SRX212649: STAT1, CD14+ monocytes | http://dbarchive.biosciencedbc.jp/kyushu-u/hg19/eachData/bw/SRX212649.bw |
| Figure S6A | SRX831874: STAT5B, CD8+ T cells | http://dbarchive.biosciencedbc.jp/kyushu-u/hg19/eachData/bw/SRX831874.bw |
| Figure S6A | SRX092314: GATA3, Th1 cells | http://dbarchive.biosciencedbc.jp/kyushu-u/hg19/eachData/bw/SRX092314.bw |
| Figure S6A | SRX831878: STAT5B, CD8+ T cells | http://dbarchive.biosciencedbc.jp/kyushu-u/hg19/eachData/bw/SRX831878.bw |
| Figure S6A | SRX1023791: SPI1, Macrophages | http://dbarchive.biosciencedbc.jp/kyushu-u/hg19/eachData/bw/SRX1023791.bw |
| Figure S6A | SRX831879: STAT5B, CD8+ T cells | http://dbarchive.biosciencedbc.jp/kyushu-u/hg19/eachData/bw/SRX831879.bw |
| Figure S6A | SRX1023790: SPI1, Macrophages | http://dbarchive.biosciencedbc.jp/kyushu-u/hg19/eachData/bw/SRX1023790.bw |
| Figure S6A | SRX831875: STAT5B, CD8+ T cells | http://dbarchive.biosciencedbc.jp/kyushu-u/hg19/eachData/bw/SRX831875.bw |
| Figure S6A | SRX150605: RELA, GM12891 | http://dbarchive.biosciencedbc.jp/kyushu-u/hg19/eachData/bw/SRX150605.bw |
| Figure S6A | SRX627428: SPI1, Dendritic cells | http://dbarchive.biosciencedbc.jp/kyushu-u/hg19/eachData/bw/SRX627428.bw |
| Figure S6A | SRX1089833: SPI1, HSCs | http://dbarchive.biosciencedbc.jp/kyushu-u/hg19/eachData/bw/SRX1089833.bw |
| Figure S6B | SRX212650: STAT1, CD14+ monocytes | http://dbarchive.biosciencedbc.jp/kyushu-u/hg19/eachData/bw/SRX212650.bw |
| Figure S6B | SRX212648: STAT1, CD14+ monocytes | http://dbarchive.biosciencedbc.jp/kyushu-u/hg19/eachData/bw/SRX212648.bw |
| Figure S6B | SRX627430: SPI1, Dendritic cells | http://dbarchive.biosciencedbc.jp/kyushu-u/hg19/eachData/bw/SRX627430.bw |
| Figure S6B | SRX1023792: SPI1, Macrophages | http://dbarchive.biosciencedbc.jp/kyushu-u/hg19/eachData/bw/SRX1023792.bw |
| Figure S6B | SRX100449: HNF4G, Hep G2 | http://dbarchive.biosciencedbc.jp/kyushu-u/hg19/eachData/bw/SRX100449.bw |
| Figure S6B | SRX100497: RXRA, Hep G2 | http://dbarchive.biosciencedbc.jp/kyushu-u/hg19/eachData/bw/SRX100497.bw |
| Figure S6B | SRX1023793: SPI1, Macrophages | http://dbarchive.biosciencedbc.jp/kyushu-u/hg19/eachData/bw/SRX1023793.bw |
| Figure S6B | SRX150572: CEBPB, Hep G2 | http://dbarchive.biosciencedbc.jp/kyushu-u/hg19/eachData/bw/SRX150572.bw |
| Figure S6B | SRX751299: MED1, Adipo-MSCs | http://dbarchive.biosciencedbc.jp/kyushu-u/hg19/eachData/bw/SRX751299.bw |
| Figure S6B | SRX100552: SP1, Hep G2 | http://dbarchive.biosciencedbc.jp/kyushu-u/hg19/eachData/bw/SRX100552.bw |
| Figure S6B | SRX813775: RELA, SGBS adipocytes | http://dbarchive.biosciencedbc.jp/kyushu-u/hg19/eachData/bw/SRX813775.bw |
| Figure S6B | SRX1023790: SPI1, Macrophages | http://dbarchive.biosciencedbc.jp/kyushu-u/hg19/eachData/bw/SRX1023790.bw |
| Figure S6B | SRX100505: HNF4A, Hep G2 | http://dbarchive.biosciencedbc.jp/kyushu-u/hg19/eachData/bw/SRX100505.bw |
| Figure S6B | SRX627428: SPI1, Dendritic cells | http://dbarchive.biosciencedbc.jp/kyushu-u/hg19/eachData/bw/SRX627428.bw |
| Figure S6B | SRX150624: EP300, Hep G2 | http://dbarchive.biosciencedbc.jp/kyushu-u/hg19/eachData/bw/SRX150624.bw |

### Oki_TableS1
